# CD99 Promotes Self-renewal in Hematopoietic Stem Cells and Leukemia Stem Cells by Regulating Protein Synthesis

**DOI:** 10.1101/2023.02.21.529419

**Authors:** Yi Huang, Eda Gozel Kapti, Yuanyuan Ji, Toby Thomas, Jebrail Dempsey, Karin Mims, Iryna Berezniuk, Liang Guo, Benjamin Kroger, Wenhuo Hu, Christopher Y. Park, Stephen S. Chung

## Abstract

Blood production is sustained by hematopoietic stem cells (HSCs), which are typically the only blood cells capable of long-term self-renewal. Acute myeloid leukemia (AML) is initiated by aberrantly self-renewing malignant stem cells termed leukemia stem cells (LSCs). HSCs exhibit and depend on low levels of protein synthesis to self-renew. However, the mechanisms by which HSCs regulate protein synthesis to maintain their capacity for self-renewal in the setting of proliferative stress and leukemogenesis remain unknown. Here we show CD99, a cell surface protein upregulated in LSCs, is required for self-renewal of proliferating HSCs and LSCs. We found that loss of CD99 in HSCs and LSCs leads to increased protein synthesis and that their self-renewal capacity can be restored by translation inhibition. These data demonstrate a functional role for CD99 in constraining protein synthesis, which may promote the clonal expansion of HSCs and LSCs that leads to AML. Furthermore, they show that similar to HSCs, LSCs depend on regulated protein synthesis.

## Introduction

Hematopoietic stem cells (HSCs) have been shown to have very low rates of protein synthesis(Signer et al., 2014), which allows them to preserve their proteome quality(Hidalgo San Jose et al., 2020). HSCs are highly sensitive to increases in protein synthesis(Chen et al., 2008a; Kharas et al., 2010; Yilmaz et al., 2006; Zhang et al., 2006), which leads to their depletion via induction of the unfolded protein response (UPR)(van Galen et al., 2014) or a tumor suppressor response(Lee et al., 2010). This may protect the clonal integrity of HSCs by promoting their depletion upon acquisition of oncogenic mutations. Because of their low protein synthesis rates, HSCs are also highly sensitive to further decreases, such as with ribosomal insufficiency(Barlow et al., 2010; Jaako et al., 2011; McGowan et al., 2011; Signer *et al*., 2014) or ribosome toxins(Palchaudhuri et al., 2016). When HSCs are subjected to proliferative stress, they exhibit increased protein synthesis leading to proteotoxic stress, which may impair their long-term function (Hidalgo San Jose *et al*., 2020; Signer *et al*., 2014; van Galen *et al*., 2014). While factors such as the 4E-binding proteins (4E-BPs)(Signer et al., 2016), angiogenin (Goncalves et al., 2016), Hsf1(Kruta et al., 2021), and Bmi1(Burgess et al., 2022) have been demonstrated to promote HSC function by constraining protein synthesis, there are likely to be other factors that preserve HSC function in the setting of proliferative stress. In particular, factors that are dynamically regulated in the context of the proliferative stress underlying leukemogenesis remain to be described.

Acute myeloid leukemia (AML) is initiated and sustained by leukemia stem cells (LSCs)(Bonnet and Dick, 1997; Lapidot et al., 1994). LSCs do not immunophenotypically resemble HSCs, but rather progenitor cells with restricted self-renewal potential such as lymphoid-primed multipotent progenitors (LMPPs)(Goardon et al., 2011) or granulocyte macrophage progenitors (GMPs)(Jamieson et al., 2004; Krivtsov et al., 2006). During transformation, these progenitors gain aberrant self-renewal to become LSCs. The precise mechanisms by which this occurs remain unknown but likely include re-activation of HSC-associated molecular and cellular programs(Eppert et al., 2011; Krivtsov *et al*., 2006; Majeti et al., 2009; Somervaille and Cleary, 2006). In contrast to HSCs, progenitor cells tolerate much higher rates of protein synthesis(Signer *et al*., 2014) and are not subject to the same constraints. Because LSCs largely resemble restricted progenitor cells but can self-renew like HSCs, it is unclear if they depend on regulated protein synthesis. Furthermore, due to a relative lack of cell surface markers that are specific for LSCs, prior studies have not tested if LSCs have lower protein synthesis rates relative to bulk AML cells and thus may have missed LSC-specific dependencies on regulated protein synthesis.

We previously identified CD99 as a cell surface marker upregulated in 86% of AML cases, also showing it is expressed at the highest levels on functionally-defined LSCs (Chung et al., 2017). Other studies have demonstrated that CD99 expression is highly enriched on LSCs (Raffel et al., 2020; Vaikari et al., 2020), as well as in AMLs with high-risk characteristics such as HSC-like features(McKeown et al., 2017; Wang et al., 2021b) and resistance to targeted therapies(Wang et al., 2021a). These studies establish CD99 as a cell surface marker allowing for improved purification of LSCs to better understand their molecular features. We previously leveraged this to show that LSCs with high CD99 expression are depleted for ribosomal protein transcripts, and that CD99 negatively regulates the Src-family kinases (SFKs), with disruption of CD99 function using monoclonal antibodies (mAbs) inducing cytotoxicity in LSCs(Chung *et al*., 2017). These data suggest that CD99 not only identifies LSCs, but that it may also function to negatively regulate protein synthesis. Here, we show that both proliferating HSCs and LSCs require CD99 to negatively regulate protein synthesis to maintain their capacity for self-renewal.

## Results

### CD99 is dispensable for steady-state hematopoiesis

To elucidate the mechanisms by which CD99 promotes self-renewal in HSCs, we evaluated hematopoietic stem and progenitor (HSPC) frequencies and blood counts in CD99-deficient mice generated by gene trapping (*B6-Cd99^Gt(pU-21T)44lmeg^*, referred to hereafter as CD99 Gt/Gt)(Park et al., 2012). We confirmed the absence of CD99 protein expression in T-cells from CD99 Gt/Gt mice by flow cytometry, given that T-cells exhibit uniformly high levels of CD99 expression (**Fig.1A**). At 10-12 weeks of age, homozygous CD99 Gt/Gt mice exhibited a significant increase in bone marrow (BM) cellularity (**Fig.1 B**), as well as a significant increase in the absolute numbers of common myeloid progenitors (CMPs; lineage-negative [LN] c-kit+sca-1+CD34+CD16/32-), granulocyte macrophage progenitors (GMPs; LN c-kit+sca-1+CD34+CD16/32+), and megakaryocyte erythroid progenitors (MEPs; LN c-kit+sca-1+CD34-CD16/32-) as compared with littermate *Cd99* wild-type (WT) mice (**Fig.1C**). However, evaluation of HSPCs by flow cytometry revealed no difference in the absolute frequencies of HSCs (LN c-kit+sca-1+CD34-CD150+), multipotent progenitors (MPPs; LN c-kit+sca-1+CD34+CD150+/-), or LN c-kit+sca-1+ cells (LSK) (**Fig.1D**). CD99 Gt/Gt mice did not demonstrate any differences in white blood cell (WBC) counts and platelet counts as compared with WT mice (**Fig.1E**). Although CD99 Gt/Gt mice had hemoglobin (Hgb) levels within normal limits, they exhibited a mild decrease as compared with WT mice (mean 14.5 g/dl, range 13.4–15.5 g/dl vs. mean 15.3 g/dl, range 14.8– 16.1 g/dl) and hematocrit (mean 45.4%, range 42.7%–49.7% vs. mean 49.4%, range 47.6%– 52.1%), along with an increase in mean corpuscular volume (MCV) (mean 46.9 fL, range 45.5– 48.5% vs. mean 45.9 fL, range 44.9–46.8%) (**Fig.1F**). To test HSPC function *in vitro*, we plated 150 purified HSCs from CD99 Gt/Gt mice and WT controls in methylcellulose semi-solid media containing myeloid-erythroid cytokines (rmIL-3, rmSCF, rh-EPO, and rh-IL6). There was no significant difference in the number of colonies derived from CD99 Gt/Gt vs. WT HSCs on initial plating, as well as upon four rounds of serial replating (**Fig.1G**). These data demonstrate that CD99 is not required for steady state hematopoiesis.

**Figure 1.**
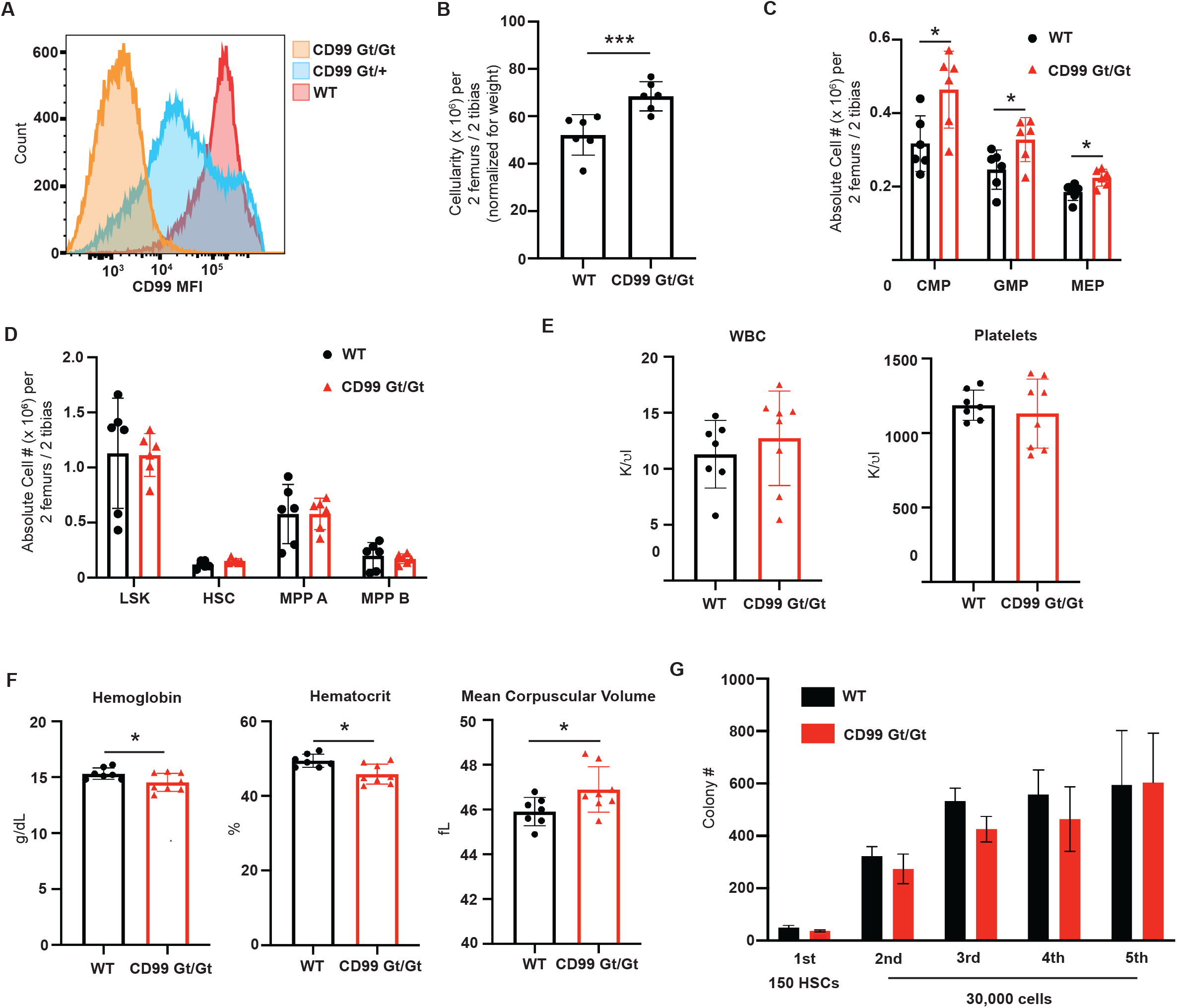
CD99 mice demonstrate no overt hematopoietic defects at steady state. (**A**) Representative flow cytometry histogram showing expression of CD99, as measured by flow cytometry on CD3+ T-cells from CD99 Gt/Gt, CD99 Gt/+, and wild-type (WT) mice. (**B**) Bone marrow (BM) cellularity of 10-12 week old CD99 Gt/Gt (n=6) and littermate WT mice (n=6). (**C**) Absolute number of common myeloid progenitors (CMP; lineage negative [LN] c-kit+sca-1-CD34+CD16/32-), granulocyte macrophage progenitors (GMP; LN c-kit+sca-1-CD34+CD16/32+), and megakaryocyte erythroid progenitors (MEP; LN c-kit+sca-1-CD34-CD16/32-) (n=6 per genotype). (**D**) Absolute number of LN c-kit+sca-1+ (LSK) cells, hematopoietic stem cells (HSC; LSK CD34-CD150+), multipotent progenitor A (MPP A; LSK CD34+CD150+), and multipotent progenitor B (MPP B; LSK CD34+CD150-) (n=6 per genotype). (**E-F**) Complete blood counts of WT and CD99 KO mice, showing white blood cell (WBC) and platelet counts, as well as hemoglobin, hematocrit, and mean corpuscular volume (n=6 per genotype). (**G**) Numbers of colonies formed 10 days after plating of 150 HSCs from three independent mice in methylcellulose containing myeloid-erythroid cytokines. Numbers of colonies 10 days after 30,000 cells were resuspended from the initial plating and replated for four successive rounds. Statistical significance was assessed using two-tailed student’s *t*-tests; *p<0.05, ***p<0.005; All data represent mean ± standard error.

### CD99 loss impairs the function of HSCs in transplantation assays

To test if loss of CD99 impairs the function of HSCs, we purified HSCs from CD99 Gt/Gt mice and WT littermate controls and transplanted them into lethally irradiated recipients along with 2 x 10^5^ unfractionated recipient BM cells (**Fig.2A**). At early time points post-transplant, peripheral blood (PB) analysis revealed similar total donor chimerism in recipients of CD99 Gt/Gt and WT HSCs. However, over 24 weeks there emerged a non-significant trend towards decreased donor chimerism in recipients of CD99 Gt/Gt HSCs (**Fig.2B**). While there was no statistically significant difference in total, B-cell, or T-cell donor chimerism in mice transplanted with CD99 Gt/Gt HSCs (**Fig.2C**), there was a significant and progressive decline in myeloid donor chimerism beginning at 12 weeks (**Fig.2D**), consistent with gradual loss of multilineage output from CD99 Gt/Gt HSCs. Evaluation of recipient BMs at 24 weeks revealed no differences in the frequency or absolute numbers of donor CD99 Gt/Gt HSCs (**Fig.2E**), but there was a significant decrease in the absolute numbers of myeloid progenitors (LN Sca-1-c-Kit+) including CMPs, GMPs, and MEPs, as well as BM cellularity in mice transplanted with CD99 Gt/Gt HSCs (**Fig.2F-G**), consistent with decreased HSC functional output. To test the self-renewal of CD99 Gt/Gt HSCs, we performed non-competitive secondary transplants using unfractionated BM from primary recipients (**Fig.2H**). Secondary recipients of BM from primary mice transplanted with CD99 Gt/Gt HSCs exhibited a marked decrease in donor chimerism in all lineages, consistent with a defect in HSC self-renewal (**Fig.2H, Supp.Fig.1A**). To exclude the possibility that our results could be due to an alteration in the cell-surface phenotype of HSCs secondary to CD99 loss, we transplanted 2 x 10^6^ unfractionated BM cells into lethally irradiated recipients. We observed no difference in total donor chimerism between recipients of BM from CD99 Gt/Gt and WT mice, and no sustained differences in donor chimerism in all lineages. However, upon secondary transplantation of 4 x 10^6^ unfractionated BM cells into lethally irradiated secondary recipients we observed in recipients of BM from CD99 Gt/Gt mice a significant decrease in total, B-cell, and T-cell donor chimerism (**Fig.2I, Supp.Fig.1B)**. Finally, to test the relative fitness of CD99 Gt/Gt HSCs, we performed competitive transplants in which 5 x 10^5^ unfractionated BM cells from CD99 Gt/Gt mice or WT controls were transplanted along with 5 x 10^5^ competitor BM cells from WT mice into lethally irradiated recipients. PB analysis revealed a significant decrease in total donor chimerism, as well as donor chimerism in all lineages (**Fig.2J, Supp.Fig.1C**). Upon secondary transplantation of 4 x 10^6^ unfractionated BM cells into lethally irradiated secondary recipients, we observed in recipients of CD99 Gt/Gt mice significantly fewer mice with multilineage re-constitution (**Fig.2K, Supp.Fig1D**). Together, these results demonstrate that CD99 is required for HSC self-renewal in the context of transplantation.

**Figure 2.**
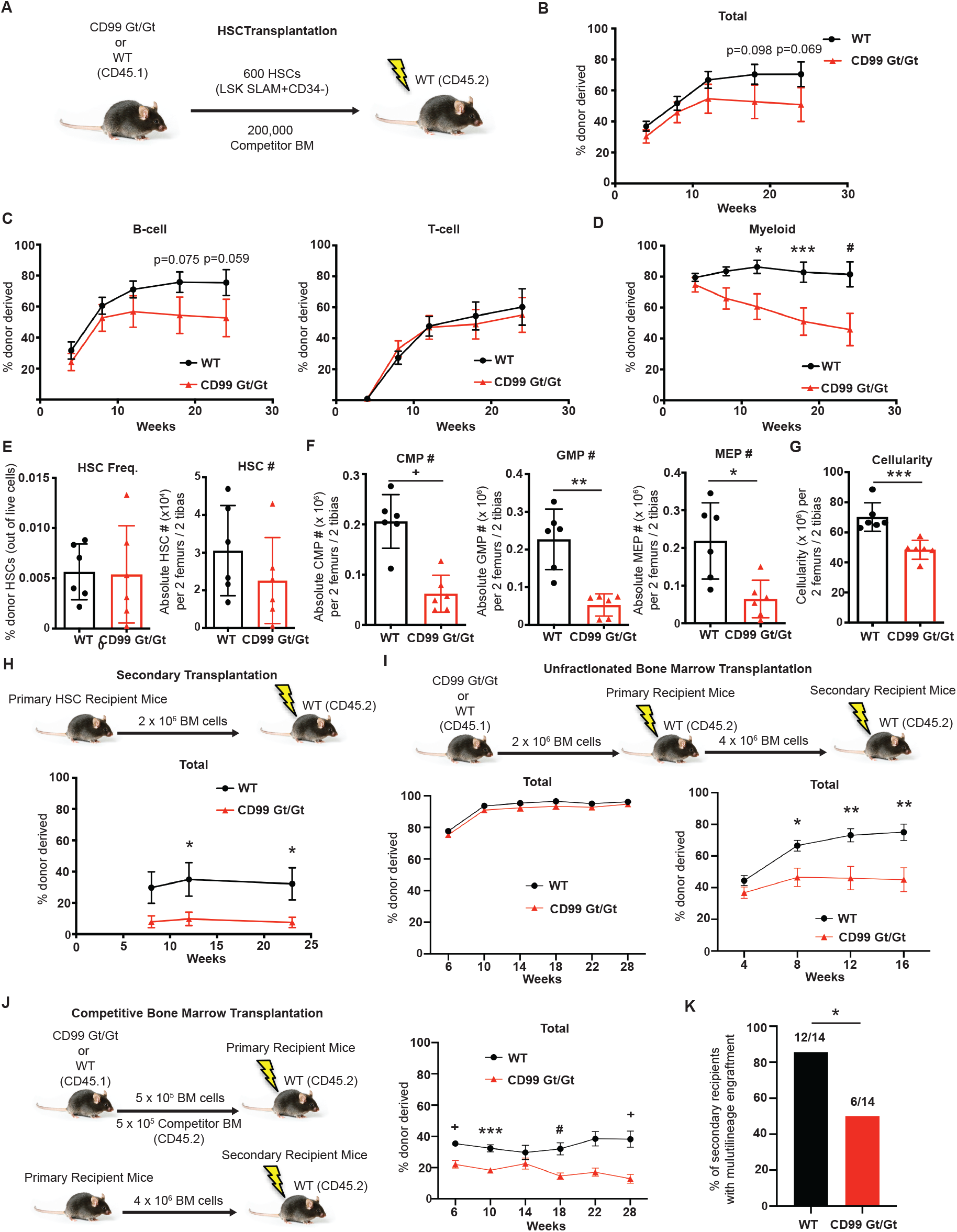
CD99 is required for HSC Self-renewal. (**A**) Schematic of primary transplants of purified HSCs. (**B**) Total donor-derived peripheral blood (PB) chimerism in primary recipients (n=6 donors and 6 recipients per genotype). (**C-D**) Donor-derived B-cell, T-cell, and myeloid cell chimerism in primary recipients (see legend above panels). (**E**) Frequency and absolute number of donor-derived HSCs in the BM of primary recipients after 24 weeks. (**F**) Absolute number of donor-derived CMPs, GMPs, and MEPs in the BM of primary recipients. (**G**) BM cellularity per two femurs and two tibias. (**H**) Schematic of secondary transplantation and total donor-derived PB chimerism in secondary recipients (n=6 donors and 6 recipients per genotype). (**I**) Schematic of primary and secondary transplants of unfractionated BM. Total donor-derived PB chimerism in primary recipients (n=6 donors and n=10 recipients per genotype) and secondary recipients (n=6 donors and n=10 recipients per genotype). (**J**) Schematic of primary and secondary competitive transplants of unfractionated BM. Total donor-derived PB chimerism in primary recipients (n=6 donors and n=10 recipients per genotype). (**K**) Number of secondary recipients with multilineage engraftment (defined as >0.5% donor myeloid and lymphoid cells) at sixteen weeks after transplant. Statistical significance was assessed using two-tailed student’s *t*-tests (**A-J**) and Fisher’s exact test (**K**); *p<0.05, **p<0.01, ***p<0.005, +p<0.0005; p-values for selected non significant trends are also shown; data represent mean ± standard error (**A-D, H-J**) and mean ± standard deviation (**E-G**).

### Loss of CD99 leads to molecular features of dysregulated protein synthesis in HSCs

To identify potential mechanisms by which CD99 promotes HSC self-renewal, we performed RNA-sequencing on HSCs purified from CD99 Gt/Gt mice and WT littermate controls. We identified 327 differentially expressed genes (p<0.01)(**Fig.3A**). Gene set enrichment analysis (GSEA) identified in CD99 Gt/Gt HSCs a decrease in expression of genes associated with HSC and LSC function, consistent with their impaired self-renewal potential (**Fig.3B**). We also observed enrichment in CD99 Gt/Gt HSCs for expression of a set of 100 genes induced in human leukemia cell lines by mAbs that disrupt the function of CD99 (Chung *et al*., 2017), suggesting a similar functional role for CD99 in mouse HSCs and human AML cells. Although we observed significant depletion of ribosomal protein transcripts, we also identified significant enrichment for transcripts associated with mTOR signaling, c-MYC targets, the proteasome, and the unfolded protein response. This led us to hypothesize that CD99 Gt/Gt HSCs may be characterized by increased protein synthesis leading to proteotoxic stress, which has been shown to promote transcriptional repression of ribosomal proteins(DuRose et al., 2009). To test if similar pathways are operative in progenitor cells, we also performed RNA-sequencing on GMPs purified from CD99 Gt/Gt mice and WT littermate controls, identifying 33 differentially expressed genes (FDR<0.05)(**Fig.3C**). Of note, only the ribosomal protein transcript and unfolded protein response signatures were concordant with our findings in HSCs, with other gene sets identified in HSCs either enriched in the opposite direction (HSC/LSC genes, MYC targets, and proteasome) or not significantly enriched (mTOR signaling and anti-CD99 mAb induced genes)(**Fig.3D**). Together, these results suggested that CD99 may function specifically in HSCs to promote self-renewal by negatively regulating protein synthesis to prevent induction of proteotoxic stress.

**Figure 3.**
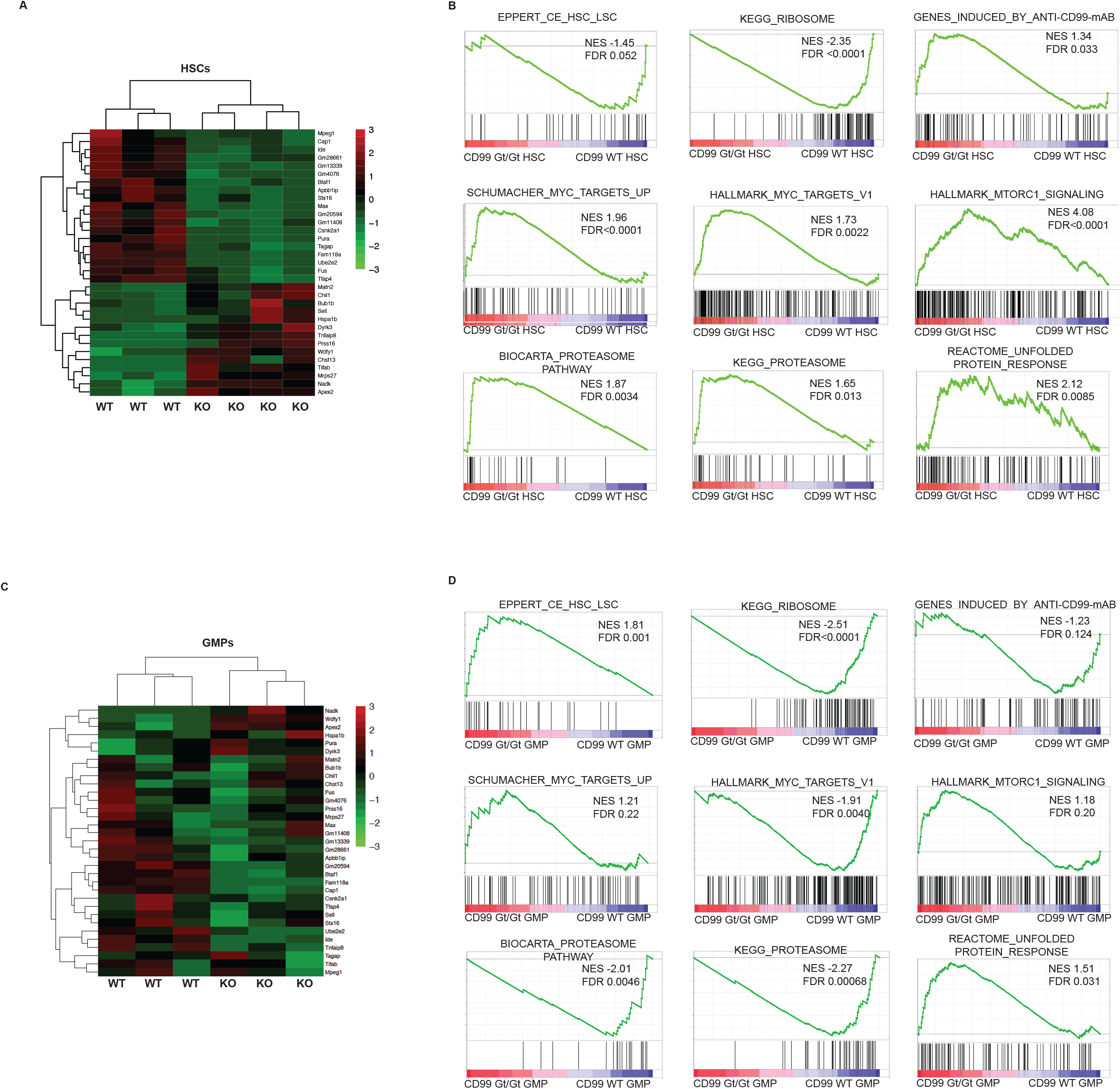
RNA-sequencing of CD99 Gt/Gt HSCs reveals features of decreased self-renewal and proteotoxic stress. (**A**) Heatmap of mRNA expression, showing genes differentially expressed between HSCs purified from CD99 Gt/Gt (KO) and wild-type littermate control (WT) mice (n=3 per genotype). (**B**) Gene set enrichment analysis (GSEA) of key biological processes enriched or depleted between KO HSCs and WT HSCs is shown. (**C**) Heatmap of mRNA expression, showing differentially expressed genes between GMPs purified from KO and WT mice. (**D**) GSEA of biological processes identified to be enriched or depleted in HSCs in KO vs. WT GMPs.

### CD99 constrains protein synthesis in proliferating HSCs to maintain proteostasis

We next sought to directly test if loss of CD99 leads to increased protein synthesis in HSCs. To do so we measured protein synthesis rates in CD99 Gt/Gt and WT HSCs *in vivo* using the puromycin analog O-propargyl-puromycin (OP-puro). OP-puro is an alkyne analog of puromycin that incorporates into nascent polypeptide chains at a rate proportional to global protein synthesis, allowing for quantification of these rates *in vivo* (Liu et al., 2012; Signer *et al*., 2014). According to published protocols(Hidalgo San Jose and Signer, 2019), we injected CD99 Gt/Gt mice and WT littermate controls with OP-puro (50 mg/kg) one hour prior to sacrifice, followed by intracellular flow cytometry to measure OP-puro incorporation in HSCs. At steady state, CD99 Gt/Gt HSCs exhibited a non-significant trend towards increased protein synthesis (p=0.063, **Fig.4A**). Given that CD99 Gt/Gt mice had minimal hematopoietic defects at steady state but CD99 Gt/Gt HSCs exhibited marked functional impairment upon transplantation, we hypothesized that CD99 becomes required for HSC self-renewal in the context of proliferative stressors. To test this, we induced HSC proliferation in CD99 Gt/Gt mice and WT controls with one dose of granulocyte colony stimulating factor (G-CSF) followed by two daily doses of cyclophosphamide (Cy)(Morrison et al., 1997), followed by protein synthesis rate quantification *in vivo* with OP-puro (**Fig.4B**). Upon G-CSF/Cy treatment, we observed a 1.68-fold increase (p=0.0001) in the cell surface expression of CD99 on HSCs in WT mice, suggesting that CD99 may be upregulated in response to proliferative stress (**Fig.4C**). We observed that HSCs undergoing cell division (S, G2, and M phases, or S/G2/M) had increased protein synthesis rates, as has been previously described(Signer *et al*., 2014). We found that CD99 Gt/Gt HSCs driven to enter the cell cycle by G-CSF/Cy exhibit a significant increase in protein synthesis compared with WT HSCs, but CD99 Gt/Gt HSCs that remain in G_0_ and G_1_ phases of the cell cycle did not (**Fig.4E**). To test if CD99 is required to constrain protein synthesis in the setting of other stressors that promote protein synthesis, we cultured CD99 Gt/Gt HSCs *ex vivo*, a context which has been demonstrated to induce a massive increase in protein synthesis (Kruta *et al*., 2021) (**Fig.4F**). After 18 hours of *ex vivo* culture, CD99 Gt/Gt HSCs in all phases of the cell cycle exhibited significantly increased protein synthesis rates compared with WT HSCs (**Fig.4G**). These data demonstrate that loss of CD99 leads to dysregulated protein synthesis in HSCs when they are subjected to stressors that increase protein synthesis, such as proliferation and *ex vivo* culture.

**Figure 4.**
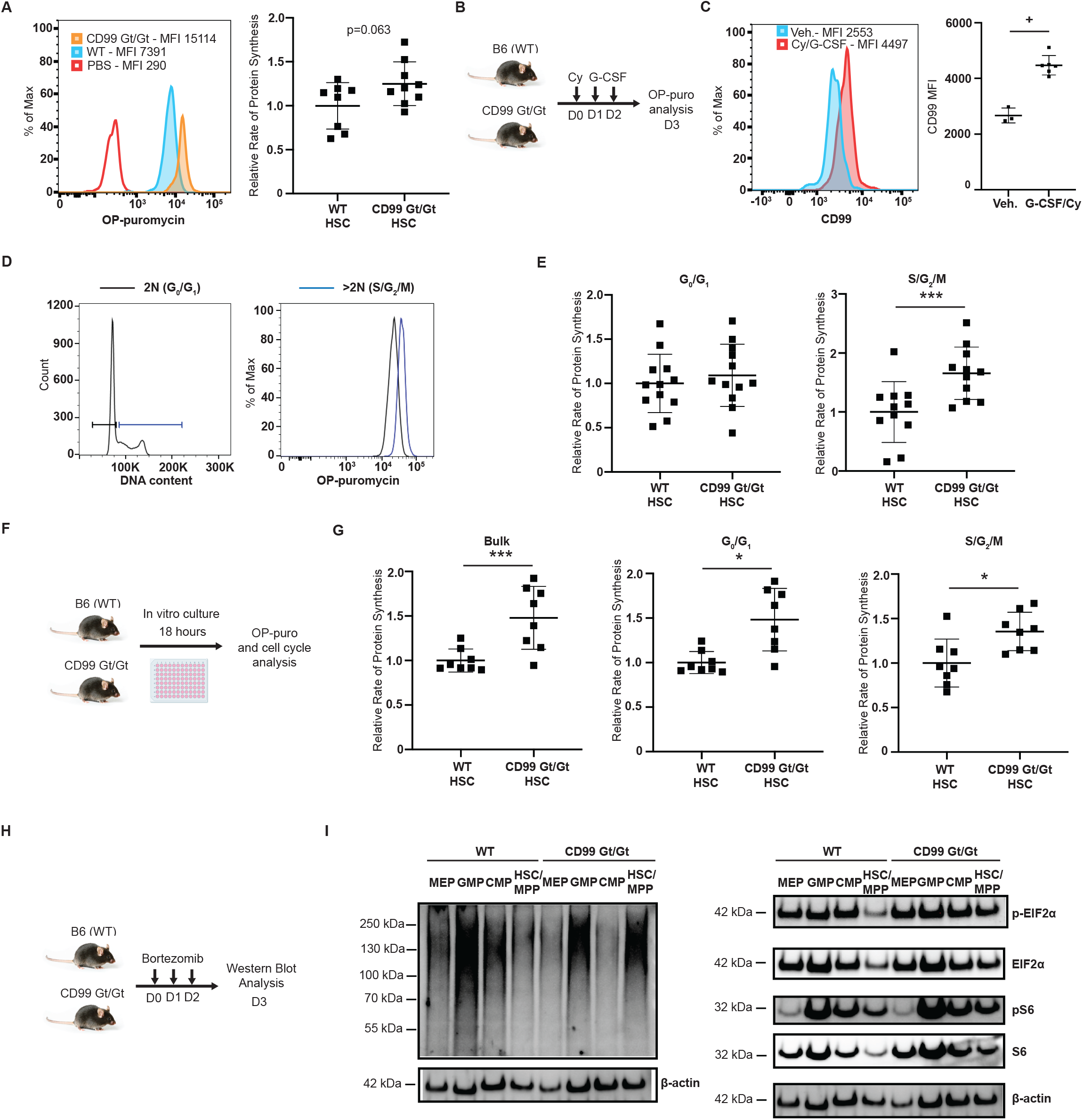
Loss of CD99 leads to increased protein synthesis and impaired proteostasis in HSCs. (**A**) *In vivo* O-propargyl-puromycin (OP-puro) incorporation in HSCs. Representative histogram with mean fluorescence intensity (MFI) indicated for CD99 Gt/Gt and WT mice injected with OP-puro and a PBS control injected mouse (left panel). *In vivo* protein synthesis in CD99 Gt/Gt and WT HSCs (n=9 and 8, respectively), normalized to the mean in WT HSCs (right panel). (**B**) Schematic for treatment of CD99 Gt/Gt and WT mice with cyclophosphamide (Cy, 4 mg) followed by two daily doses of G-CSF (5 μg). (**C**) CD99 cell surface protein expression on WT HSCs following Cy/G-CSF treatment. Representative histogram is shown (left panel). CD99 MFI for vehicle (Veh.) and Cy/G-CSF treated mice (n=3 and 6, respectively). (**D**) Representative gating strategy based on DNA content to identify 2N (G_0_/G_1_) and >2N (S/G2/M) cells. (**E**) *In vivo* protein synthesis in CD99 Gt/Gt and WT HSCs (n=12 per genotype) following Cy/G-CSF treatment relative to the mean in WT HSCs for cells in G_0_/G_1_ (left panel) and S/G2/M (right panel). (**F**) Schematic for *ex vivo* culture of HSCs followed by OP-puro analysis. (**G**) Protein synthesis in *ex vivo* cultured CD99 Gt/Gt and WT HSCs (n=12 per genotype), normalized to the mean in WT HSCs for total cells (left panel), as well as those in G_0_/G_1_ (middle panel) and S/G2/M (right panel). (**H**) Schematic for treatment of CD99 Gt/Gt and WT mice with three daily doses of bortezomib (1 mg/kg) followed by western blot analysis of hematopoietic stem and progenitor cells. (**I**) Western blots examining ubiquitylated protein, p-eIF2a, eIF2α, pS6, S6, and ß-actin in 3 x 10^4^ sorted HSC/MPPs, CMPs, GMPs, and MEPs from CD99 Gt/Gt and WT mice treated with bortezomib. Statistical significance was assessed using two-tailed student’s *t*-tests; *p<0.05, ***p<0.005, +p<0.0005; p-values for selected non-significant trends are also shown; data represent mean ± standard deviation.

To determine if increased protein synthesis in CD99 Gt/Gt HSCs leads to impaired proteostasis, we evaluated ubiquitylated protein levels and activation of the integrated stress response (ISR) by western blot. We examined phosphorylated eIF2α (p-eIF2a) as a marker of the ISR(Kimball et al., 1998; Krishnamoorthy et al., 2001), as well as phosphorylated S6 (p-S6) as a marker for increased mTORC1 signaling. We did not find any significant differences in these measurements between CD99 Gt/Gt and WT control HSCs/MPPs (LSK CD48-) in steady state conditions (**Supp.Fig.2A**), consistent with the absence of significant hematologic abnormalities in this state. When we induced HSCs to proliferate with G-CSF/Cy, we also did not find any significant differences between CD99 Gt/Gt and WT control HSCs/MPPs (**Supp.Fig.2B**). We reasoned that the short three-day time frame of the experiment, as well as the dilution of misfolded proteins enabled by cell division, may limit the effects of CD99 loss on proteostasis. Thus, to increase the accumulation of misfolded proteins we treated mice with the proteasome inhibitor bortezomib (1 mg/kg) for three consecutive days (**Fig.4H**). CD99 Gt/Gt HSCs/MPPs exhibited increased ubiquitylated protein and p-EIF2a as compared with WT control HSCs/MPPs, consistent with the accumulation of misfolded proteins and activation of the ISR (**Fig.4I**). We also observed increased p-S6 in CD99 Gt/Gt HSCs/MPPs, suggesting that upon induction of proteotoxic stress and activation of the ISR, they fail to decrease protein synthesis downstream of dysregulated mTORC1 signaling. Therefore, the function of CD99 in negatively regulating protein synthesis allows HSCs/MPPs to avoid accumulation of misfolded proteins and activation of the ISR.

### Rapamycin rescues the self-renewal defect of CD99 Gt/Gt HSCs

To test if increased protein synthesis is responsible for the impaired self-renewal of proliferating CD99 Gt/Gt HSCs, we assessed whether inhibition of translation could restore their capacity for self-renewal. Given our data suggestive of increased mTORC1 signaling in CD99 Gt/Gt HSCs, such as enrichment for mTORC1 signaling associated genes (**Fig.3B**) and increased S6 phosphorylation (**Fig.4I**), we chose to inhibit protein synthesis using the mTORC1 inhibitor rapamycin. Furthermore, rapamycin has been demonstrated to rescue loss of HSC self-renewal in the setting of increased mTOR signaling(Chen et al., 2009), such as with deletion of *Pten*(Yilmaz *et al*., 2006) and *Pml*(Bernardi et al., 2006), and mTORC2 signaling has been shown to be dispensable for the function of adult HSCs(Magee et al., 2012). We repeated transplants of HSCs purified from CD99 Gt/Gt mice and WT littermate controls into lethally irradiated recipient mice. Beginning 48 hours after transplantation, mice were injected with rapamycin (4 mg/kg, intraperitoneally) daily for two weeks, followed by every other day continuously (**Fig.5A**). At 24 weeks after transplant, vehicle treated recipients of CD99 Gt/Gt HSCs demonstrated a significant decrease in total PB donor chimerism as compared with vehicle treated recipients of WT HSCs (**Fig.5B**). Treatment of recipients of CD99 Gt/Gt HSCs with rapamycin led to a non-significant trend towards increased engraftment. Conversely, rapamycin treated recipients of WT HSCs exhibited a significant decrease in engraftment compared with vehicle treated recipients WT HSCs, consistent with the sensitivity of HSCs to further decreases in protein synthesis(Cai et al., 2015; Palchaudhuri *et al*., 2016; Signer *et al*., 2014). We observed the same differences in myeloid donor chimerism, where rapamycin additionally led to a statistically significant increase in the engraftment of CD99 Gt/Gt HSCs (**Fig.5C**). We also observed similar trends in T-cell and B-cell donor chimerism (**Supp.Fig.3A-C**). Evaluation of recipient BMs at 24 weeks revealed a significant decrease in donor HSC frequency in vehicle treated recipients transplanted with CD99 Gt/Gt HSCs as compared with WT HSCs (**Fig.5D**). Treatment of recipients of CD99 Gt/Gt HSCs with rapamycin led to a complete rescue of this decrease in HSC frequency. Bone marrow cellularity did not differ across genotypes and treatment groups (**Supp.Fig.3D**). The same pattern of decreased donor cell engraftment in vehicle treated recipients of CD99 Gt/Gt HSCs that was rescued with rapamycin treatment was seen when evaluating absolute numbers and frequencies of donor HSCs, GMPs, CMPs, and MEPs (**Fig.5E**, **Supp.Fig.3E**). These data demonstrate that the decreased number and functional output of CD99 Gt/Gt HSCs can be rescued by inhibition of mTORC1 signaling.

**Figure 5.**
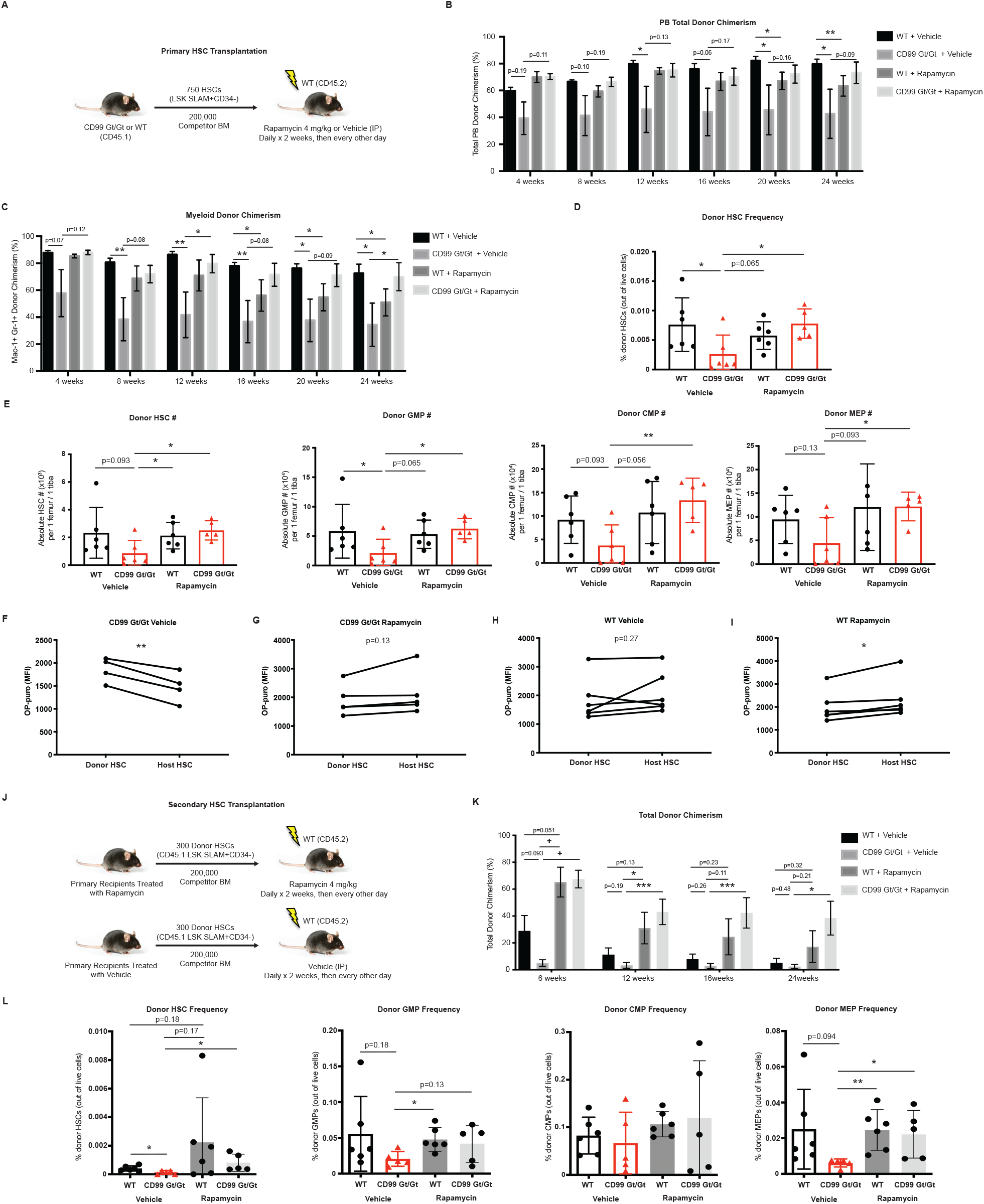
Rapamycin rescues the self-renewal defect of CD99 Gt/Gt HSCs. (**A**) Schematic of primary transplants of purified HSCs from CD99 Gt/Gt mice and WT littermate controls followed by treatment of recipient mice with rapamycin. (**B**) Total donor-derived PB chimerism in primary recipients (n=6 donors and 6 recipients per genotype) over the course of 24 weeks. (**C**) Donor-derived myeloid chimerism in the PB over the course of 24 weeks. (**D**) Frequency of donor-derived HSCs in the BM of primary recipients after 24 weeks. (**E**) Absolute number of donor-derived HSCs, GMPs, CMPs, and MEPs in the BM of primary recipients after 24 weeks. (**F-I**) *Ex vivo* OP-puro incorporation in donor (CD45.1) vs. host (CD45.2) HSCs isolated from the BM of recipients at 24 weeks (see labels above panels for experimental groups). (**J**) Schematic of secondary transplants of HSCs purified from primary recipients into secondary recipients treated with rapamycin or vehicle control. (**K**) Total donor-derived PB chimerism in secondary recipients (n=6 donors and 6 recipients per genotype) over 24 weeks. (**L**) Frequency of donor-derived HSCs, GMPs, CMPs, and MEPs in the BM of secondary recipients after 24 weeks. Statistical significance was assessed using two-tailed student’s *t*-tests (**B-C,F-I,K**) and Mann-Whitney *U* tests (**E,L**); *p<0.05, **p<0.01, ***p<0.005, +p<0.0005; p-values for selected non-significant trends are also shown; data represent mean ± standard error (**B-C, K**) and mean ± standard deviation (**E,L**).

To test the effects of CD99 loss and rapamycin treatment on protein synthesis in HSCs in the context of transplantation, we performed *ex vivo* OP-puro incorporation assays on HSCs at the end of primary transplant experiments. We furthermore compared protein synthesis rates in donor vs. host HSCs in each recipient. In a pairwise analysis, we found that in vehicle treated recipients of CD99 Gt/Gt HSCs, donor HSCs exhibited a significant increase in protein synthesis rates compared with host HSCs (**Fig.5F**), consistent with our prior observation that CD99 Gt/Gt HSCs driven to proliferate have increased protein synthesis rates (**Fig.4E**). Treatment with rapamycin completely abrogated these increased protein synthesis rates in donor HSCs (**Fig.5G**), consistent with the functional rescue of CD99 Gt/Gt HSCs that we observed in this experimental group. In contrast, in vehicle treated recipients of WT HSCs, donor HSCs did not exhibit any differences in protein synthesis rates compared with host HSCs (**Fig.5H**). Treatment with rapamycin led to a decrease in protein synthesis rates in donor HSCs as compared with host HSCs(**Fig.5I**), consistent with their decreased function and a genotype-dependent difference in response to rapamycin (**Fig.5B-C**). Thus, the decreased function of CD99 Gt/Gt HSCs is due to increased protein synthesis that can be rescued with mTORC1 inhibition, consistent with the need for HSCs to maintain protein synthesis at a low homeostatic set point. The decreased function of WT HSCs in response to mTORC1 inhibition is consistent with the sensitivity of HSCs to further decreases in their already low levels of protein synthesis(Barlow *et al*., 2010; Jaako *et al*., 2011; McGowan *et al*., 2011; Palchaudhuri *et al*., 2016; Signer *et al*., 2014)

Finally, to formally test whether the self-renewal defect of CD99 Gt/Gt HSCs can be rescued by inhibition of protein synthesis, we performed secondary transplant experiments. Given the decreased HSC-frequency in primary recipients of CD99 Gt/Gt HSCs, we purified equal numbers of donor HSCs from primary recipients and transplanted them into lethally irradiated secondary recipients, who were then continued on either rapamycin or vehicle as in the corresponding primary recipients (**Fig.5J**). At 24 weeks after transplant, total donor PB chimerism was lowest in recipients of CD99 Gt/Gt HSCs, with complete rescue of their engraftment levels with rapamycin treatment (**Fig.5K**). A similar rescue phenotype was observed in myeloid cells, B-cells, and T-cells (**Supp.Fig.4A-B**), consistent with restoration of multilineage output from CD99 Gt/Gt HSCs. At early time points after transplantation, rapamycin also increased donor engraftment from WT HSCs (**Fig.5K**, **Supp.Fig.4A-B**), but this effect was not sustained. Evaluation of recipient BMs at 24 weeks revealed a significant decrease in donor HSC frequency in vehicle treated recipients transplanted with CD99 Gt/Gt HSCs as compared with WT HSCs (**Fig.5L**). Treatment of recipients of CD99 Gt/Gt HSCs with rapamycin again led to a complete rescue of this decrease in HSC frequency. Similar results were obtained in independent experiments where three million unfractionated BM cells were transplanted from primary recipients into secondary recipients (**Supp.Fig.4C-D**). Thus, the self-renewal defect associated with loss of CD99 is due to increased protein synthesis downstream of mTORC1 signaling.

### CD99 promotes LSC function in AML1-ETO-induced leukemia by regulating protein synthesis

Given that CD99 is required on HSCs undergoing proliferative stress and CD99 is enriched in the vast majority of human LSCs (Chung *et al*., 2017), we hypothesized that LSCs may upregulate CD99 in response to the proliferative stress of leukemogenesis. We thus sought to determine whether CD99 promotes LSC self-renewal by negatively regulating protein synthesis. To identify subtypes of AML most likely to be dependent on CD99, we assessed CD99 transcript expression on 398 patients enrolled on the Eastern Cooperative Group (ECOG) E1900 clinical trial and correlated CD99 expression with the presence of recurrent AML mutations (**Fig.6A**). We found that AMLs harboring *AML1-ETO* translocations exhibited among the highest levels of CD99 expression (Somers’ D correlation metric 0.45 compared with other subtypes). We thus utilized a well-established retro-viral transduction-transplantation model of leukemogenesis driven by an alternatively spliced isoform of the *AML1-ETO* transcript, *AML1-ETO9a(Yan et al., 2006*). HSPC-enriched c-kit positive cells were isolated from the BM of CD99 Gt/Gt mice or WT littermate controls treated with 5-FU (200 mg/kg), followed by retroviral transduction to express *AML1-ETO9a* and green fluorescent protein (GFP) from a bi-cistronic vector (**Fig.6B**). Transduced GFP-positive cells were transplanted into lethally irradiated WT recipients along with 2 x 10^5^ unfractionated recipient BM cells. To test whether dysregulated protein synthesis contributed to any observed phenotypes, we further treated recipient mice with rapamycin or vehicle control. Mice were monitored for disease development and overall survival. The efficiency of leukemogenesis and survival of primary transplanted mice varied significantly within experimental groups, consistent with the described long latency of these leukemias the need for stochastically arising cooperating mutations(Hatlen et al., 2016) (**Fig.6C**). The leukemias that developed lacked lineage marker expression and sca-1 expression, with c-kit expression on a subset of cells, as has been described(Yan *et al*., 2006), and we did not observe any immunophenotypic differences between CD99 Gt/Gt and WT-derived leukemias (**Fig.6D**).

**Figure 6.**
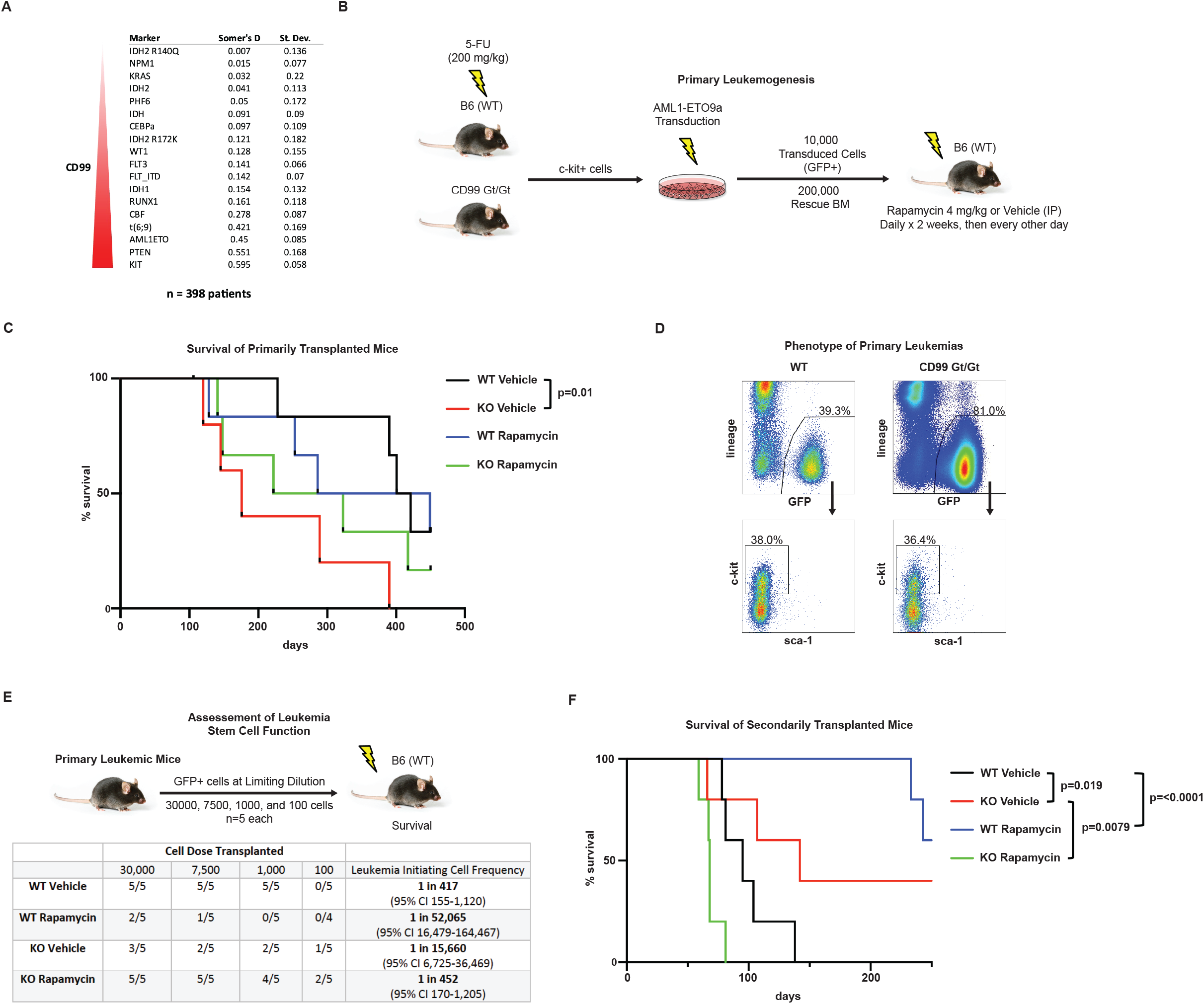
Loss of CD99 leads to an impairment in LSC function that can be rescued with rapamycin treatment. (**A**) CD99 transcript levels were assessed in 398 AML cases from the ECOG1900 clinical trial and correlated with the presence of recurrent cytogenetic abnormalities and mutations (Somers’ D test). The top 15 CD99-associated genetic abnormalities (out of 36 tested) are shown. (**B**) Schematic of transduction of c-kit positive HSPCs from 5-fluorouracil (5-FU) treated CD99 Gt/Gt mice and WT littermate controls to co-express *AML1-ETO9a* and green fluorescent protein (GFP), followed by transplant into lethally irradiated primary recipients. Beginning 48 hours after transplant, recipients were treated with rapamycin or vehicle control at the indicated dose and frequency. (**C**) Overall survival of mice transplanted with transduced CD99 Gt/Gt or WT HSPCs and treated with vehicle or rapamycin (KO Vehicle, WT Vehicle, KO Rapamycin, and WT Rapamycin, respectively). (n=6 for each group except for n=5 in the KO Vehicle group). (**D**) Immunophenotype of GFP-positive leukemia cells derived from WT and CD99 Gt/Gt mice. (**E**) Schematic of secondary transplant of GFP-positive leukemia cells that developed in primary recipients into secondary recipient mice at limiting dilution. The numbers of mice that engrafted with and succumbed to leukemia out of the numbers transplanted with each cell number are indicated in the table. Leukemia-initiating cell frequency is shown as calculated using Poisson statistics. (**F**) Overall survival of mice secondarily transplanted with KO Vehicle, WT Vehicle, KO Rapamycin, and WT Rapamycin leukemias at the 30,000, 7,500, and 1,000 cell doses (n=15 for each group). Statistical significance was assessed using Log-rank (Mantel-Cox) tests.

To quantify functional LSCs in these leukemias, we performed transplants of primary leukemia cells at limiting dilution into secondary recipients (**Fig.6E**). Due to the direct antileukemic effects that rapamycin may have on fully transformed leukemia cells (Recher et al., 2005), we did not treat secondary recipients with rapamycin. CD99 Gt/Gt-derived leukemias that were generated in vehicle treated primary recipients (KO Vehicle) had a much lower LSC frequency (1 in 15,660, 95% CI 6,725-36,469) as compared with WT-derived leukemias generated in vehicle treated primary recipients (WT Vehicle)(1 in 417, 95% CI 155-1,120), consistent with loss of CD99 leading to an impairment in LSC function. CD99 Gt/Gt-derived leukemias generated in rapamycin treated primary recipients (KO Rapamycin) exhibited a complete restoration of LSC frequency (1 in 452, 95% CI 170-1,205) to levels comparable to WT Vehicle leukemias. Conversely, WT-derived leukemias generated in rapamycin treated primary recipients (WT Rapamycin) exhibited a marked decrease in LSC-frequency (1 in 52,065, 95% CI 16,479-164,467) compared with WT Vehicle leukemias (**Fig.6E**). Thus, rapamycin increases LSC frequency in CD99 Gt/Gt leukemias but decreases LSC frequency in WT leukemias. Secondary recipients of KO Vehicle leukemias exhibited improved survival (median not reached, range 61-156 days) compared with secondary recipients of WT Vehicle leukemias (median of 114 days, range 77-156 days)(p=0.019)(**Fig.6F**). Secondary recipients of KO Rapamycin leukemias had a significant decrease in survival (median 68 days, range 59-117 days) as compared with recipients of KO Vehicle leukemias (p=0.0079), consistent with rescue of LSC function with rapamycin treatment. In contrast, secondary recipients of WT Rapamycin leukemias had a significant increase in survival (median not reached) compared with recipients of WT Vehicle leukemias (p<0.0001), consistent with loss of LSC function. Thus, loss of CD99 impairs LSC function in *AML1-ETO*9a-driven leukemia, and inhibition of protein synthesis with rapamycin can completely rescue this defect. Conversely, rapamycin impairs the function of WT LSCs, suggesting that similar to HSCs, LSCs may be characterized by low protein synthesis rates and highly sensitive to further decreases.

To test if CD99 Gt/Gt-derived leukemias exhibit increased protein synthesis, as well as to assess the effects of rapamycin on protein synthesis during leukemogenesis, we measured protein synthesis rates in *AML1-ETO*9a-driven primary and secondarily transplanted leukemias using *ex vivo* OP-puro incorporation assays. Although the immunophenotype of LSCs in this model has not been well defined, we examined c-kit positive cells to further focus these studies on a population of cells likely enriched for LSCs. KO Vehicle leukemias exhibited increased protein synthesis rates as compared with WT Vehicle leukemias (**Fig.7A**). KO Rapamycin leukemias had lower protein synthesis rates compared with KO Vehicle leukemias, suggesting that rapamycin allowed for the persistence and selection for LSCs with lower protein synthesis rates. Conversely, WT Rapamycin leukemias had higher protein synthesis rates, suggesting that rapamycin promoted the persistence and selection for LSCs with higher protein synthesis rates. Notably, the experimental groups with the lowest protein synthesis rates correlated with those with the highest LSC frequencies as tested in **Fig. 6**. These differences in protein synthesis rates persisted through secondary transplants in the absence of rapamycin treatment (**Fig.7B**), suggesting that such differences that developed during leukemogenesis were inherent cell autonomous properties of the resulting leukemias. Additionally, we observed similar differences in protein synthesis rates in bulk leukemia cell populations (**Supp.Fig.5A-B**), consistent with the notion that properties of LSCs may be propagated to bulk leukemia cells(Eppert *et al*., 2011; Majeti *et al*., 2009). Western blot analysis revealed increased pS6 in KO Vehicle and WT Rapamycin leukemias, consistent with the increased protein synthesis rates we observed in these experimental groups (**Fig.7C**). We observed increased ubiquitylated protein in KO Vehicle leukemias (**Fig.7D**), suggesting that leukemias lacking CD99 accumulate more misfolded proteins, but we did not observe activation of the ISR (**Supp.Fig.5C**), and treatment with rapamycin did not appear to decrease the levels of these ubiquitylated proteins (**Fig.7D**), suggesting that other pathways may mediate the loss of self-renewal that characterizes these LSCs. Finally, RNA-sequencing of leukemias revealed that rapamycin promoted a marked depletion of ribosomal protein transcripts in CD99 Gt/Gt leukemias but not WT leukemias (**Fig.7E**), consistent with the genotype-dependent differential response of these leukemias to rapamycin. Together, these results demonstrate that loss of CD99 leads to increased protein synthesis rates in LSCs, and reduction of protein synthesis rates with rapamycin has opposing effects on CD99 Gt/Gt and WT LSCs (**Fig.7F**). This establishes a model in which LSCs are similar to HSCs and require protein synthesis rates to be tightly regulated at a low homeostatic set-point.

**Figure 7.**
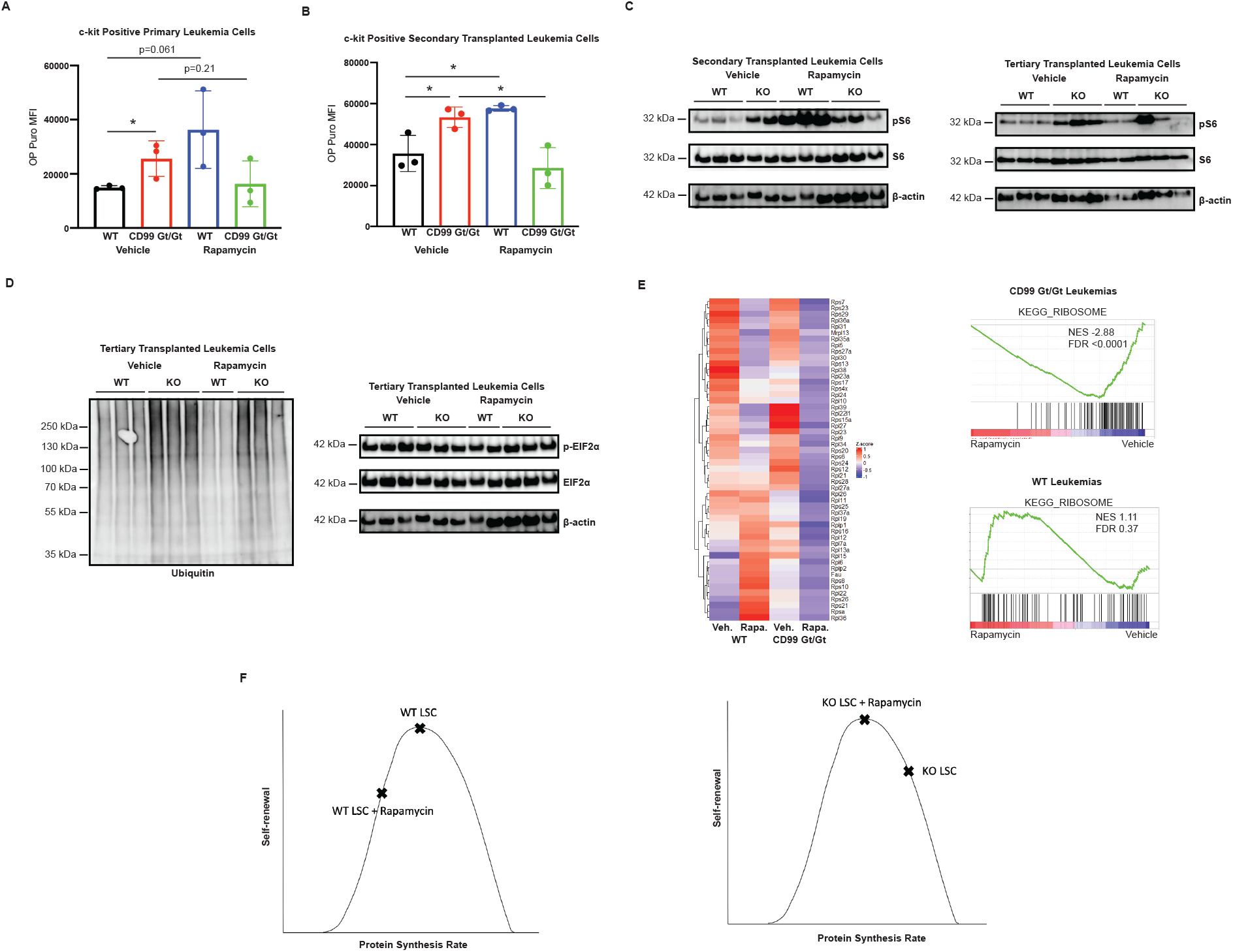
Loss of CD99 leads to increased protein synthesis in AML-ETO-driven leukemias. (**A-B**) *Ex vivo* OP-puro incorporation in c-kit positive leukemia cells from primary or secondary recipients of CD99 Gt/Gt or WT HSPCs treated with vehicle or rapamycin (KO Vehicle, WT Vehicle, KO Rapamycin, and WT Rapamycin, respectively). (**C-D**) Western blots examining pS6, S6, ß-actin, ubiquitylated protein, p-eIF2a, and eIF2α in 3 x 10^4^ bulk leukemia cells from each experimental group (derived from secondary or tertiary transplanted mice as indicated). (**E**) Heatmap and GSEA analysis depicting ribosomal protein transcripts differentially expressed in the four experimental groups (n=3 biological replicates per group). (**F**) Model of differential response of CD99 Gt/Gt and WT LSCs to rapamycin. Statistical significance was assessed using two-tailed student’s *t*-tests; *p<0.05; p-values for selected non-significant trends are also shown; data represent mean ± standard deviation.

## Discussion

Low protein synthesis rates have been shown to characterize somatic stem cells in many tissues(Blanco et al., 2016; Llorens-Bobadilla et al., 2015; Sampath et al., 2008; Sanchez et al., 2016; Zismanov et al., 2016), including the hematopoietic system(Signer *et al*., 2014). In HSCs, increases in protein synthesis likely lead to their depletion via induction of a tumor suppressor response(Lee *et al*., 2010) and the ISR(van Galen *et al*., 2014; van Galen et al., 2018). However, the mechanisms that allow HSCs to avoid loss of self-renewal during physiologic processes that require them to expand remain unclear. Such mechanisms may also be relevant to pathologic conditions, such as the pre-leukemic clonal expansion that precedes the development of AML(Jan et al., 2012; Shlush et al., 2014).

Because LSCs adopt a restricted progenitor-like immunophenotype(Goardon *et al*., 2011; Krivtsov *et al*., 2006) but can self-renew like HSCs, it remains unresolved if LSCs behave more like HSCs or restricted progenitors with respect to their dependence upon regulated protein synthesis. LSCs are thought to aberrantly self-renew by adopting features of HSCs(Krivtsov *et al*., 2006), but they must also undergo a massive clonal expansions during leukemogenesis that likely require increases in protein synthesis. It is thus possible that they depend on similar mechanisms as proliferating HSCs to maintain their capacity for self-renewal in the face of increased protein synthesis. Prior studies have shown that *Pten*-inactivation leads to depletion of HSCs and development of leukemia in mice(Yilmaz *et al*., 2006; Zhang *et al*., 2006), suggesting that LSCs can tolerate high protein synthesis rates, and that they accomplish this by partially inactivating other tumor suppressors(Lee *et al*., 2010). However, not all AMLs inactivate tumor suppressors(Papaemmanuil et al., 2016), and other models that promote increases in protein synthesis (Chen *et al*., 2008a) do not develop leukemia. Additionally, due to a lack of LSC-specific cell surface markers, prior studies have not determined if LSCs have lower protein synthesis rates relative to bulk AML cells.

In our studies we show that a specific marker of LSCs, CD99(Chung *et al*., 2017), is dispensable for steady state hematopoiesis but becomes required for HSC self-renewal under conditions that require HSCs to proliferate, such as with transplantation or chemotherapeutic stress. We show that CD99 is upregulated on HSCs in the setting of such proliferative stressors, and that it promotes self-renewal by negatively regulating protein synthesis. Reasoning that leukemogenesis represents another proliferative stressor on stem cells in the hematopoietic system, we additionally show that CD99 plays a similar functional role in LSCs. Importantly, we observed opposing effects of protein synthesis inhibition based on CD99 genotype in both HSCs and LSCs. This is consistent with the well described observation that HSCs require tight regulation of protein synthesis and are sensitive to both increases and decreases in protein synthesis(Barlow *et al*., 2010; Chen *et al*., 2008a; Jaako *et al*., 2011; Kharas *et al*., 2010; McGowan *et al*., 2011; Palchaudhuri *et al*., 2016; Signer *et al*., 2014; Yilmaz *et al*., 2006; Zhang *et al*., 2006). By demonstrating a similar function for CD99 in HSCs and LSCs, these results strongly suggest that LSCs co-opt from HSCs a dependence on tightly regulated protein synthesis to self-renew. Moreover, although other LSC-specific markers such as GPR56(Pabst et al., 2016; Raffel *et al*., 2020) have been described to enrich for LSCs among AML cells, our studies are unique in identifying a direct functional role for an LSC marker in regulation of self-renewal.

We acknowledge several limitations to these studies. First, although rapamycin inhibits protein synthesis, its effects on mTORC1 signaling may affect many other cellular pathways such as autophagy and biosynthetic processes. Further studies using pharmacologic inhibitors and genetic models that directly inhibit translation initiation and/or ribosomal biogenesis may shed light on the contribution of other processes regulated downstream of mTORC1. Nevertheless, the tight correlation between protein synthesis rates and the self-renewal phenotypes and molecular features we observed strongly suggest that the predominant effect of rapamycin in our studies is on protein synthesis. Second, retrovirally-driven leukemia models are limited by poor control of and often supraphysiologic levels of oncogene expression(Chen et al., 2008b), as well as artifacts from *ex vivo* culture during transduction. Future experiments testing the effects of CD99 loss in genetically faithful models of AML will be needed to overcome these limitations.

Our findings lay the groundwork for future studies to better understand the mechanisms by which proliferating HSCs and LSCs maintain their capacity for self-renewal by regulating protein synthesis rates. We speculate that during pre-leukemic clonal expansion of HSCs, driver mutations that activate certain signaling pathways that induce greater increases in protein synthesis may be more dependent on CD99. Future studies using genetic models of clonal hematopoiesis, such as with loss of *Dnmt3a* or *Tet2*, promise to define the role of CD99 in this process. Studies using different genetic models of AML will also be important to define the breadth of genetic sub-types of AML that require CD99 for LSC self-renewal. Additional work is also needed to determine the pathways that lead to depletion of LSCs upon dysregulation of protein synthesis, and whether they are the same as or unique from those induced in HSCs. Finally, alterations in signaling (e.g. mTOR, MAPK) that decrease global protein synthesis paradoxically increase the translational efficiency of specific mRNAs, particularly in the context of stress and/or transformed phenotypes(Blanco *et al*., 2016; Rao et al., 2012; Zismanov *et al*., 2016). Parsimonious translation of such mRNAs has been shown in some stem cells to be essential for self-renewal(Blanco *et al*., 2016; Sampath *et al*., 2008; Zismanov *et al*., 2016). Thus, negative regulation of global protein synthesis by CD99 may lead to paradoxically increased translation of important regulators of LSC function, and studies of selective mRNA translation, such as with ribosome profiling(Ingolia, 2016; Ingolia et al., 2012), promise to reveal novel regulators of LSC function.

In summary, we have identified a novel function for CD99 in regulating protein synthesis in proliferating HSCs and LSCs to promote their self-renewal. Our work provides a physiologically relevant model in which to study the effects of perturbations in global protein synthesis on stem cell function. Our results also reveal regulated protein synthesis as a fundamental property of stem cells that is co-opted by LSCs, setting the stage for future studies to test whether it is also a therapeutic liability.

## Experimental Methods

### Flow cytometry and cell sorting

Flow cytometry and fluorescence-activated cell sorting was performed on a FACSAria II cell sorter (BD Biosciences, San Jose, California). Analysis and isolation of mouse hematopoietic cells was performed using the following antibodies: CD3 (145-2C11), CD4 (RM4-5), CD8 (53-6.7), Ter119 (TER-119), Gr-1 (RB6-8C5), CD11b (M1/70), B220 (RA3-6B2), c-kit (2B8), sca-1 (D7), CD150 (TC15-12F12.2), CD16/32 (93) and CD34 (HM34) from Biolegend. A lineage stain consisting of CD3, CD4, CD8, Ter119, Gr-1, CD11b, and B220 was used. Dead cells were excluded from analyses and cell sorting using propidium iodide (0.1 μg/ml). Cells stained with CD34 antibodies were incubated for 90 minutes on ice, with all other incubations approximately 30 minutes on ice.

### Bone marrow cell isolation

Bone marrow (BM) cells were isolated for analysis by flushing two femurs and two tibias. For HSC-isolation, cells were isolated by crushing the spine, femurs, tibias, and pelvic bones of mice with a mortar and pestle into phosphate-buffered saline (PBS) without Ca^2+^ and Mg^2+^ with 2% fetal bovine serum (FBS, Sigma, St. Louis, Missouri), followed by filtration through a 70 μm strainer. Pre-enrichment for c-kit+ cells was achieved using anti-c-kit magnetic beads (Miltenyi Biotec).

### Transplantation assays

Recipient C57/Bl6 (CD45.2) mice were lethally irradiated with two 540 cGy doses (XRAD 320 irradiator, Precision X-Ray) given at least three hours apart. Within 24 hours of irradiation, the indicated donor cells were transplanted via retro-orbital sinus. For all transplants performed with purified HSCs and transduced HSPCs, 2 x 10^5^ unfractionated recipient BM cells were co-transplanted. For non-competitive BM transplants 2 x 10^6^ donor cells were transplanted and for competitive BM transplants 5 x 10^5^ donor cells and 5 x 10^5^ recipient cells were transplanted. For secondary transplants of unfractionated BM, 2-4 x 10^6^ cells pooled from 3-6 primary recipients with total donor chimerism close to the mean of each genotype were transplanted.

### Peripheral blood analysis

Blood was collected by lateral tail vein collection using capillary blood collection tubes (Greiner Bio-One MiniCollect). Automated peripheral blood counts were obtained using a Hemavet 950FS Hematology System (IDEXX) according to standard manufacturer’s instructions. In all transplant experiments, donor peripheral blood (PB) chimerism was measured in recipient mice every four weeks. PB specimens underwent cell lysis with ammonium chloride potassium buffer followed by staining with antibodies against CD45.1, CD45.2, Gr-1, Mac-1, B220, and CD3.

### Bone marrow analysis

At 24 weeks after transplant, BM cells were isolated from two femurs and two tibias of recipient mice as detailed above and stained with antibodies against CD45.1, CD45.1, lineage (CD3, CD4, CD8, B220, CD11b, Gr-1, Ter119), c-kit, sca-1, CD34, CD150, and CD16/32).

### In vitro colony-forming assays

HSCs from the BM of CD99 Gt/Gt and littermate WT mice were double sorted to >95% purity and seeded at a density of 150 cells/replicate into cytokine-supplemented methylcellulose medium (Methocult M3434; Stem Cell Technologies, Vancouver, Canada). Colonies number was scored after 10 days. Cells were resuspended in PBS and a portion was taken for replating (30,000 cells/replicate) for a total of five platings.

### Mice

All mice were housed in University of Texas Southwestern (UTSW) or Memorial Sloan Kettering Cancer Center (MSKCC) animal facilities. All animal procedures were conducted in accordance with the Guidelines for the Care and Use of Laboratory Animals and were approved by the Institutional Animal Care and Use Committees (IACUCs) at UTSW and MSKCC. B6-*Cd99^Gt(pU-21T)44lmeg^* mutant mice have been described previously (Park *et al*., 2012) and were backcrossed for more than eight generations onto a C57Bl/6 background carrying the *Ptprc^a^* allele (CD45.1), and C57Bl/6 mice carrying the *Ptprc^b^* allele (CD45.2) were used as recipient mice in transplant experiments. Both male and female mice were used in all studies.

### Administration of Rapamycin

Rapamycin (LC Laboratories, Woburn, Massachusetts) was resuspended in ethanol at 10 mg/ml and then diluted to 5% Tween-80 (Sigma) and 5% PEG-400 (Hampton Research, Aliso Viejo, California). Starting at 48 hours after transplant, mice were injected with a dose of 4 mg/kg IP daily for two weeks, followed by every other day continuously until mice were sacrificed for analysis at 24 weeks.

### RNA sequencing

HSCs, GMPs, or GFP-positive leukemia cells were sorted into Trizol LS (Thermo Fisher, Rockford, Illinois). cDNA libraries were prepared using a SMARTer mRNA amplification kit (Clontech, Mountain View, California) and sequencing was performed using the Hi-Seq platform (Illumina, San Diego, California) with 40 million paired end reads per sample.

### In vivo protein synthesis assay

OP-puro (Thermo Fisher, Rockford, Illinois) was reconstituted in PBS at a concentration of 5 mg/ml (pH 6.4-6.6). Mice were injected intraperitoneally (IP) (50 mg/kg) with OP-puro or PBS control. One hour after injection, mice were euthanized and BM was isolated and enriched for c-kit positive cells as detailed above. Cells were stained with antibodies to identify HSCs (LSK CD34-CD150+), followed by washing with PBS and fixation/permeabilization using the BD Cytofix/Cytoperm kit according to the manufacturer’s protocol (BD Biosciences, San Jose, California). Cells were then pelleted and resuspended in the Click-iT OP-puro reaction cocktail with AlexaFluor 488 picolyl azide (Thermo Fisher) at 5 μM final concentration and incubated for 30 minutes at room temperature. For specimens in which DNA content was to be measured, 4’,6-diamidino-2-phenylindole (DAPI) was added at a concentration of 10 μg/ml for the final ten minutes of the incubation. Cells were then washed twice with PBS and resuspended in PBS for analysis by flow cytometry.

### In vitro protein synthesis assay

For experiments in **Fig.4**, HSCs from CD99 Gt/Gt and WT mice were sorted into 96-well plates and cultured in DMEM (Gibco, Waltham, Massachusetts) supplemented with 200 uM L-cysteine, 50 uM 2-Mercaptoethanol, 1mM L-glutamine and 0.1% BSA. Cells were incubated at 37°C, 5% CO_2_ for 18 hours, at which point OP-puro was added to the medium (20uM final concentration) and incubated at 37°C, 5% CO_2_ for 30 minutes. Cells were then stained with antibodies to identify HSCs (LSK CD34-CD150+) and fixed/permabilized for OP-puro incorporation analysis as described above for the *in vivo* assay. For experiments in **Fig.5**, BM was isolated from primary recipient mice and c-kit+ cells were enriched as detailed above. These cells were incubated in DMEM/F12 (Gibco) supplemented with 10% FBS (Sigma), as well as stem cell factor (10ng/ul), interleukin-3 (10ng/ul), interleukin-6 (10ng/ul), thrombopoietin (10ng/ul), and ligand for fms-like tyrosine kinase receptor-3 (10ng/ul) (all from Peprotech, Cranbury, New Jersey). OP-puro was immediately added to the medium (20uM final concentration) and incubated at 37°C, 5% CO_2_ for 30 minutes. Cells were then stained with antibodies to identify HSCs and fixed/permabilized for OP-puro incorporation analysis as described above.

### Western blot analysis

The indicated numbers of HSC/MPPs, CMPs, GMPs, and MEPs were sorted directly into 25% trichloroacetic acid (TCA, Sigma). Distilled water was added to dilute the final TCA concentration to 10%, and lysates were centrifuged at 14,000 g x 15 minutes at 4°C. Precipitated protein was washed with acetone twice, allowed to dry, and resuspended in solubilization buffer (9M urea, 2% Triton X-100, 1% DTT). 4x LDS sample buffer (Thermo) was added and samples were heated at 70°C for 10 minutes prior to separation on NuPAGE Novex 4-12% Bis-Tris Protein Gels (Thermo). Western blotting was performed using the following antibodies: ubiquitin (P4D1), p-EIF2a (Ser51), EIF2α (9722), pS6 (Ser 235/236), S6 (5G10), and β-actin (13E5) from Cell Signaling Technology (Danvers, Massachusetts). Electrochemiluminescent reagent (EMD Millipore, Billerica, Massachusetts) was applied to Western blots and images were acquired using a ChemiDoc MP instrument (Biorad, Hercules, California). Densitometry analysis was performed using ImageJ analysis software.

### Retroviral transduction

CD99 Gt/Gt and WT mice were injected with 200 mg/kg 5-fluorouracil IP seven days before sacrifice, followed by isolation of c-kit positive HSPCs from the BM as described above. HSPCs were cultured overnight in RPMI (Gibco) supplemented with 10% FBS (Sigma), as well as stem cell factor (10ng/ul), interleukin-3 (10ng/ul), interleukin-6 (10ng/ul), thrombopoietin (10ng/ul), and ligand for fms-like tyrosine kinase receptor-3 (10ng/ul) (Peprotech), followed by two rounds of spinoculation (2,530rpm, 37°C for 90 minutes) at 24 and 48 hours with concentrated retrovirus in the same media supplemented with 20 μg/ml polybrene (Sigma-Aldrich). Transduction efficiency was determined by reporter GFP fluorescence at 72 hours after by flow cytometry. GFP positive cells were sorted and transplanted into lethally irradiated recipient mice with 2 x 10^5^ unfractionated recipient BM cells.

### RNA sequencing

Primary AML specimens were FACS-sorted into Trizol LS (Thermo Fisher, Rockford, Illinois). cDNA libraries were prepared using a SMARTer mRNA amplification kit (Clontech, Mountain View, California) and sequencing was performed using the Hi-Seq platform (Illumina, San Diego, California) with 40 million paired end reads per sample.

## Statistical and data analysis

All flow cytometry data were analyzed using Flowjo (TreeStar, Ashland, Oregon). L-IC frequency estimations were calculated using a Poisson distribution probability calculator (L-Calc, http://www.stemcell.com/en/Products/All-Products/LCalc-Software.aspx, Stem Cell Technologies, Vancouver, Canada). RNA-seq data was aligned by STAR(Dobin et al., 2013) to human genome hg37 and reads for each gene were counted by HTSeq(Anders et al., 2015). Differentially expressed genes were identified by DESeq2(Love et al., 2014), followed by gene set enrichment analyses(Subramanian et al., 2005). P-values were calculated using two-tailed t-tests (Prism 9, GraphPad Inc., La Jolla, California). Estimated variation was taken into account for each group of data and is indicated as standard error or standard deviation in each figure legend.

## Supplementary Materials

**Supplemental Figure 1.**
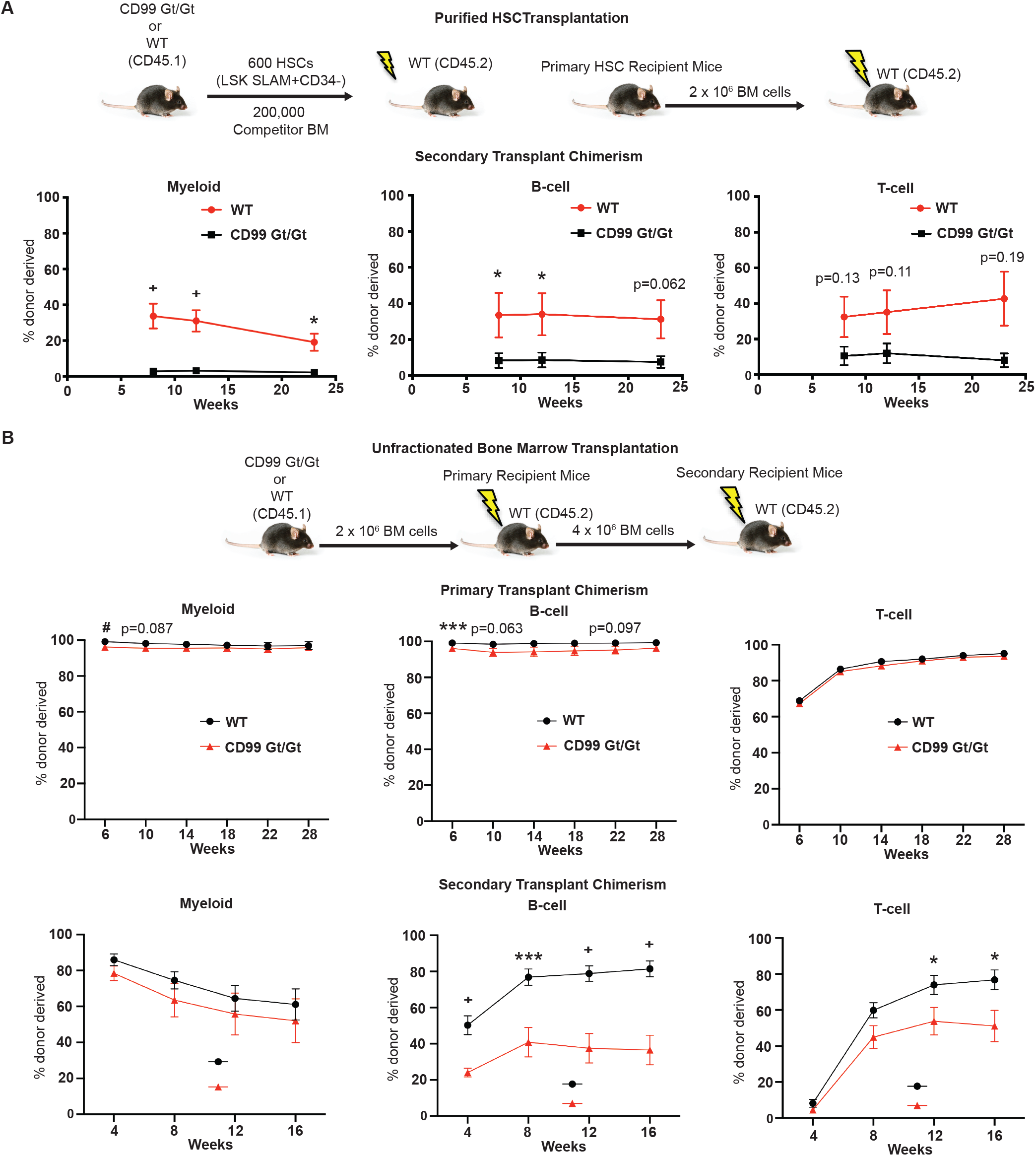
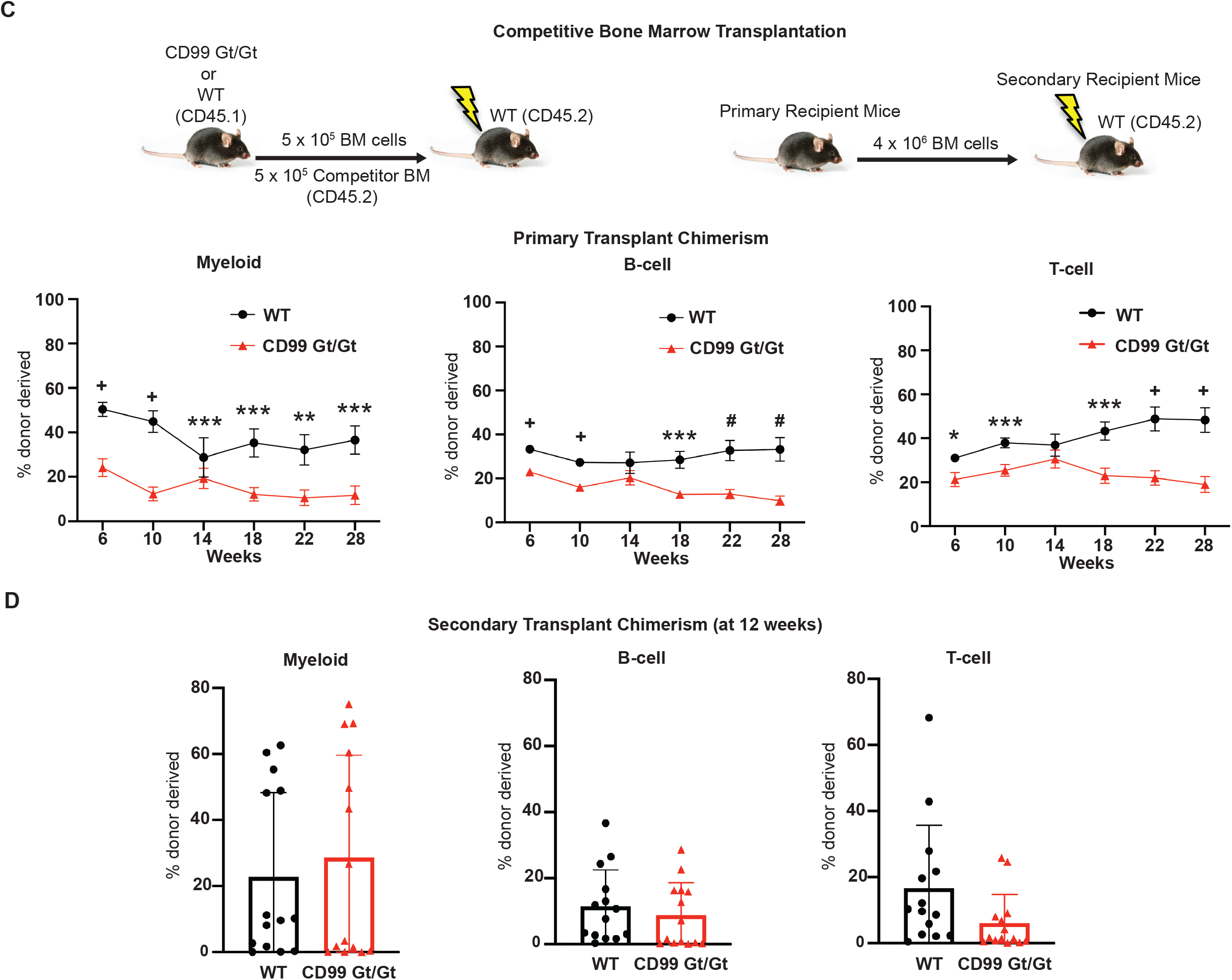
CD99 is required for HSC self-renewal. (**A**) Schematic of primary transplants of purified HSCs and secondary transplants of unfractionated BM (n=6 donors and n=6 recipients per genotype). Donor-derived myeloid, B-cell, and T-cell chimerism in primary recipients (see legend above panels). (**B**) Schematic of primary and secondary transplants of unfractionated bone marrow (BM) (n=6 donors and n=10 recipients per genotype). Donor-derived myeloid, B-cell, and T-cell chimerism in primary and secondary recipients (see legend above panels). (**C-D**) Schematic of primary and secondary competitive transplants (n=6 donors and n=10 recipients per genotype). Donor-derived myeloid, B-cell, and T-cell peripheral blood (PB) chimerism in primary and secondary recipients (see legend above panels). Statistical significance was assessed using a two-tailed student’s *t*-tests); *p<0.05, **p<0.01, ***p<0.005, #p<0.001, +p<0.0005; p-values for selected non-significant trends are shown; data represent mean ± standard error (**A-C**) and mean ± standard deviation (**D**).

**Supplemental Figure 2.**
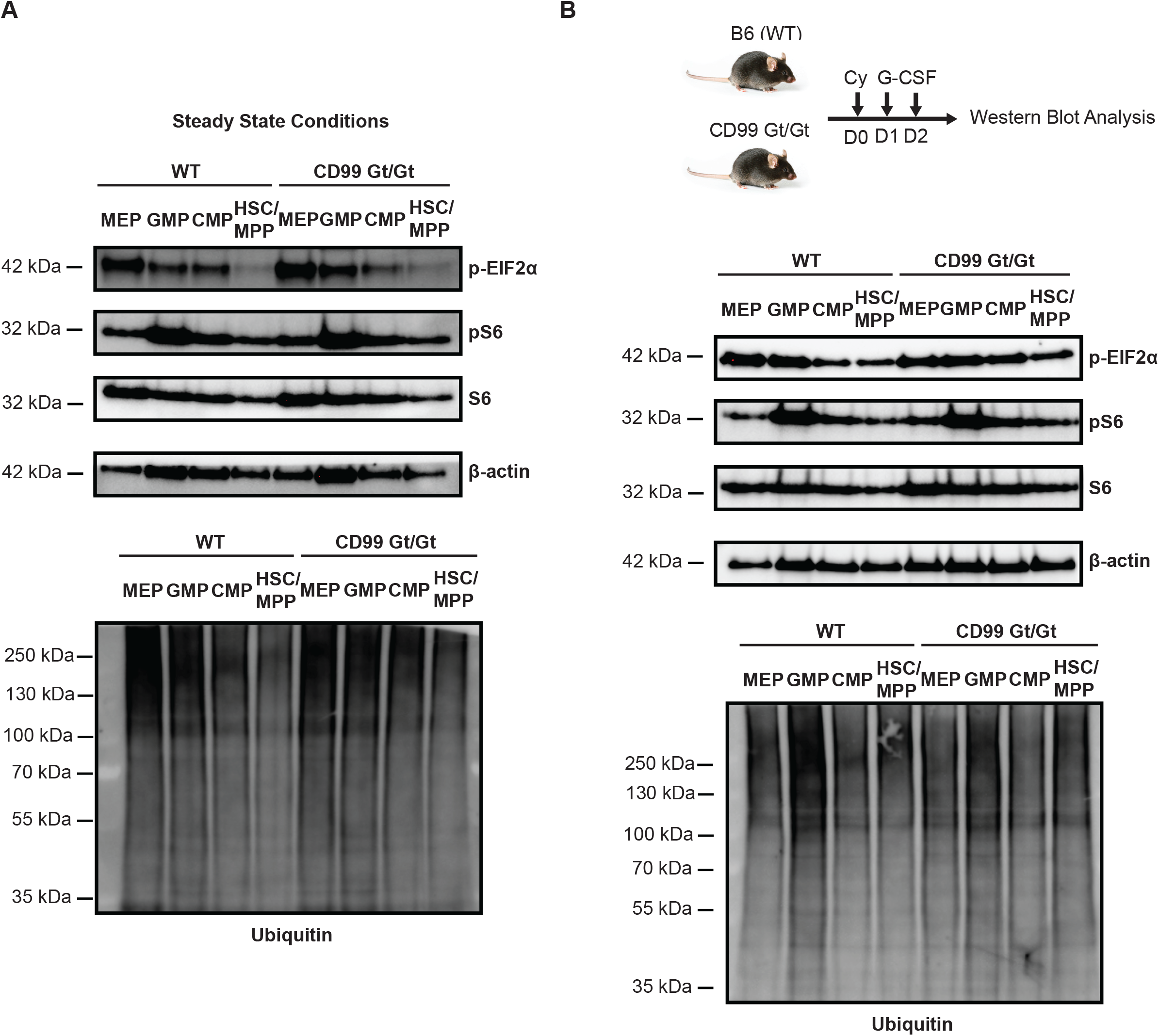
Effect of CD99 loss on protein ubiquitylation, S6 phosphorylation, and activation of the integrated stress response at steady state or with Cy/G-CSF treatment. Western blots examining p-eIF2a, eIF2a, pS6, S6, ß-actin, and ubiquitylated protein in 3 x 10^4^ HSC/MPPs, CMPs, GMPs, and MEPs from CD99 Gt/Gt and WT mice at steady state (**A**) or after treatment with Cy/G-CSF (**B**).

**Supplemental Figure 3.**
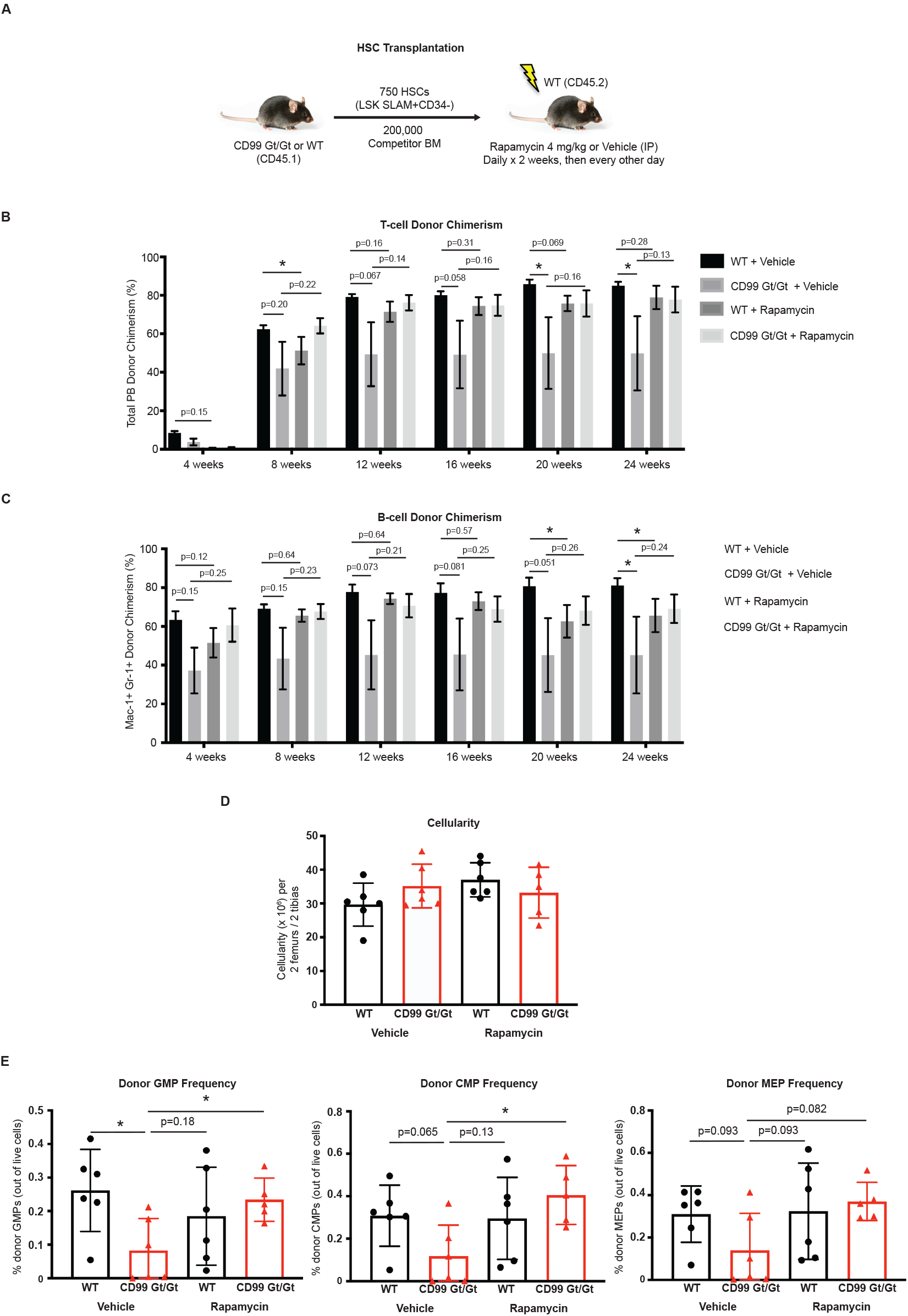
Effect of rapamycin on engraftment of CD99 Gt/Gt HSCs in primary recipients. (**A**) Schematic of primary transplants of purified HSCs from CD99 Gt/Gt mice and WT littermate controls followed by treatment of recipient mice with rapamycin or vehicle control. (**B-C**) Total donor-derived T-cell and B-cell PB chimerism in primary recipients (n=6 donors and 6 recipients per genotype) over the course of 24 weeks. (**D**) BM cellularity of primary recipients per two femurs and two tibias after 24 weeks. (**E**) Frequency of donor-derived GMPs, CMPs, and MEPs in the BM of primary recipients after 24 weeks. Statistical significance was assessed using two-tailed student’s t-tests (**B-C**) and Mann-Whitney *U* tests (**D-E**); *p<0.05; p-values for selected non-significant trends are also shown; data represent mean ± standard error (**B-C**) and mean ± standard deviation (**D-E**).

**Supplemental Figure 4.**
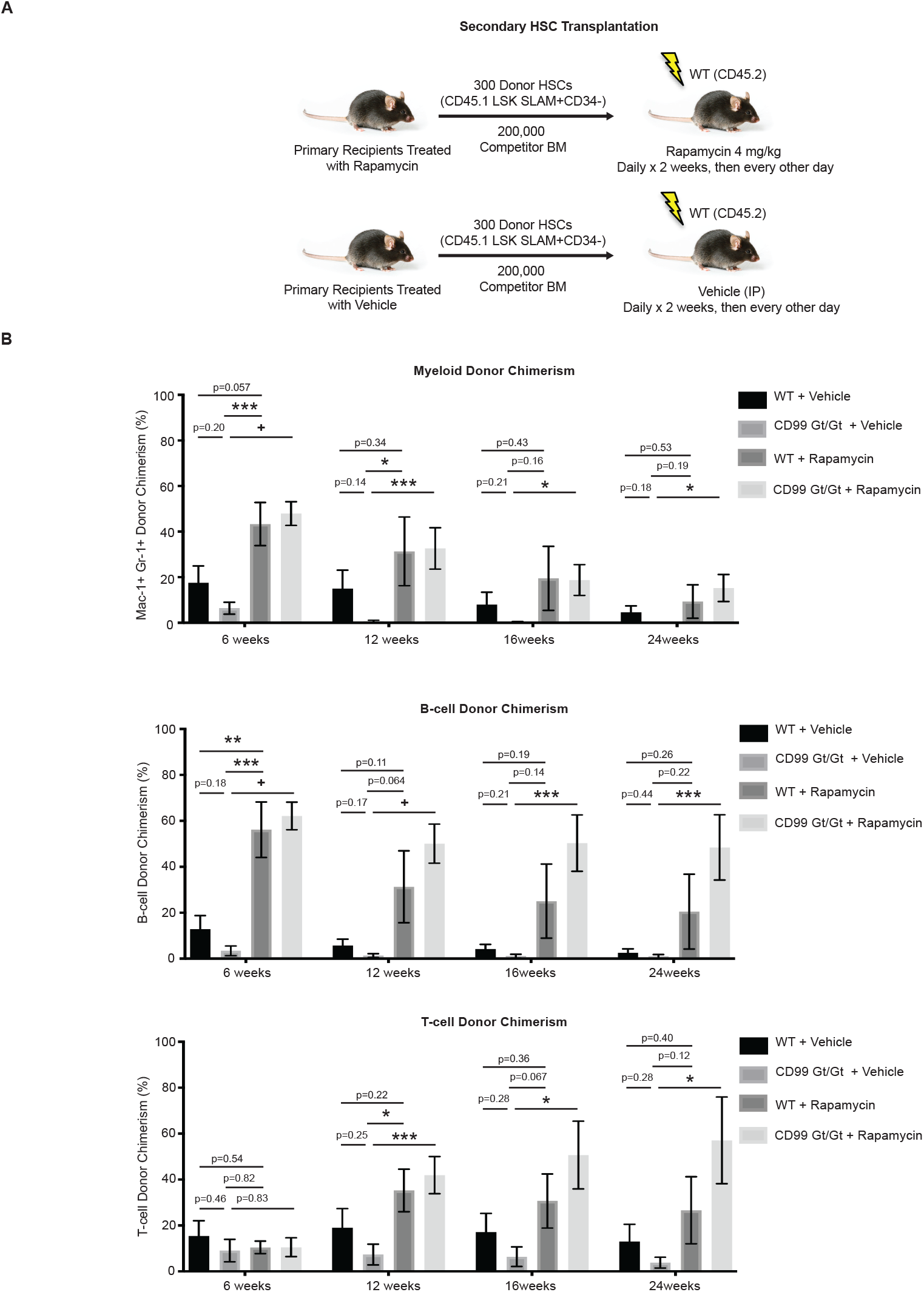
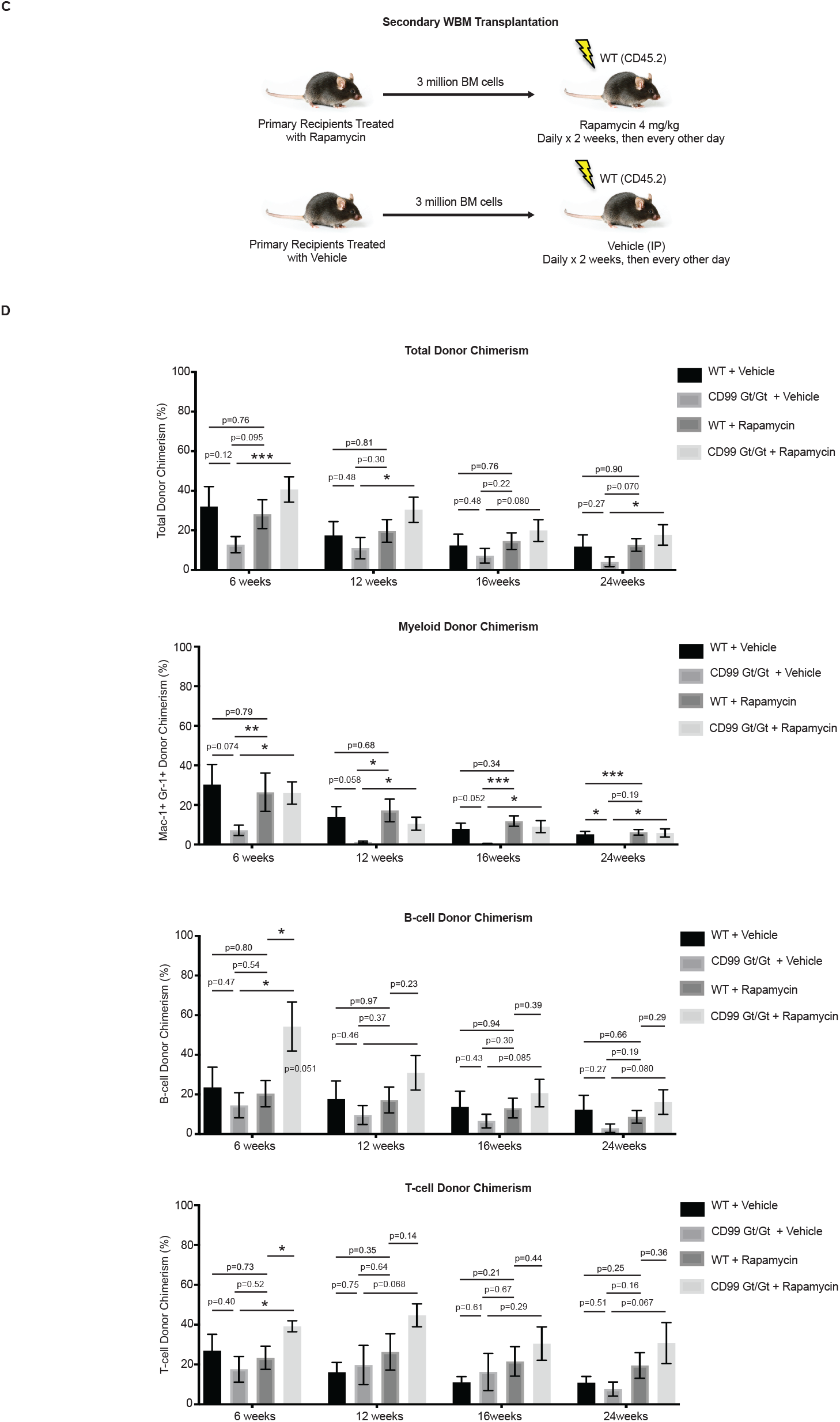
Effect of rapamycin on engraftment of CD99 Gt/Gt HSCs in secondary recipients. (**A**) Schematic of secondary transplants of donor HSCs purified from primary recipients of CD99 Gt/Gt HSCs and WT controls followed by treatment of recipient mice with rapamycin or vehicle control. (**B**) Donor-derived myeloid, T-cell, and B-cell PB chimerism in secondary recipients (n=6 donors and 6 recipients per genotype) over the course of 24 weeks. (**C**) Schematic of secondary transplants of unfractionated BM from primary recipients of CD99 Gt/Gt HSCs and WT controls followed by treatment of recipient mice with rapamycin or vehicle control. (**D**) Total donor-derived, myeloid, T-cell, and B-cell PB chimerism in secondary recipients (n=6 donors and 6 recipients per genotype) over the course of 24 weeks. Statistical significance was assessed using two-tailed student’s t-tests; *p<0.05, **p<0.01, ***p<0.005, +p<0.0005; p-values for selected non-significant trends are also shown; data represent mean ± standard error.

**Supplemental Figure 5.**
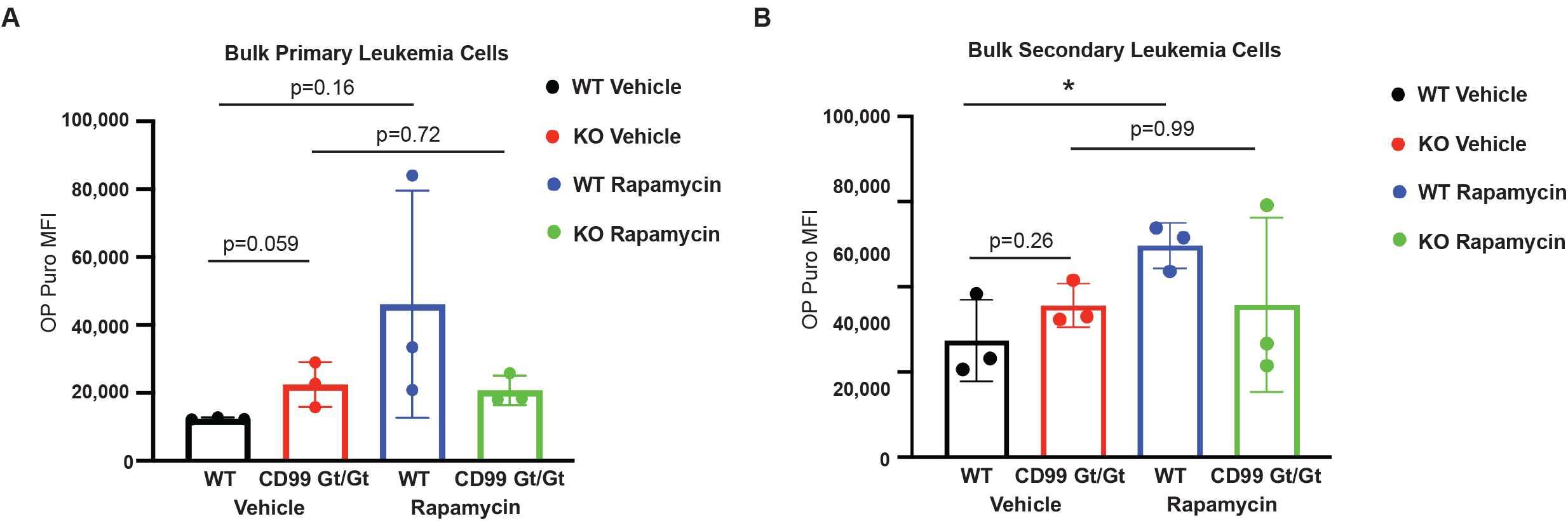
Loss of CD99 leads to increased protein synthesis in AML-ETO-driven leukemias. *Ex vivo* OP-puro incorporation in bulk leukemia cells from primary (**A**) or secondary (**B**) recipients of *AML1-ETO*9a-transduced CD99 Gt/Gt or WT HSPCs treated with vehicle or rapamycin (KO Vehicle, WT Vehicle, KO Rapamycin, and WT Rapamycin, respectively).

## References and Notes

Anders, S., Pyl, P.T., and Huber, W. (2015). HTSeq--a Python framework to work with high-throughput sequencing data. Bioinformatics 31, 166–169. 10.1093/bioinformatics/btu638.

Barlow, J.L., Drynan, L.F., Hewett, D.R., Holmes, L.R., Lorenzo-Abalde, S., Lane, A.L., Jolin, H.E., Pannell, R., Middleton, A.J., Wong, S.H., et al. (2010). A p53-dependent mechanism underlies macrocytic anemia in a mouse model of human 5q-syndrome. Nat Med 16, 59–66. nm.2063 [pii] 10.1038/nm.2063.

Bernardi, R., Guernah, I., Jin, D., Grisendi, S., Alimonti, A., Teruya-Feldstein, J., Cordon-Cardo, C., Simon, M.C., Rafii, S., and Pandolfi, P.P. (2006). PML inhibits HIF-1alpha translation and neoangiogenesis through repression of mTOR. Nature 442, 779–785. 10.1038/nature05029.

Blanco, S., Bandiera, R., Popis, M., Hussain, S., Lombard, P., Aleksic, J., Sajini, A., Tanna, H., Cortes-Garrido, R., Gkatza, N., et al. (2016). Stem cell function and stress response are controlled by protein synthesis. Nature 534, 335–340. 10.1038/nature18282.

Bonnet, D., and Dick, J.E. (1997). Human acute myeloid leukemia is organized as a hierarchy that originates from a primitive hematopoietic cell. Nat Med 3, 730–737.

Burgess, R.J., Zhao, Z., Nakada, D., and Morrison, S.J. (2022). Bmi1 suppresses protein synthesis and promotes proteostasis in hematopoietic stem cells. Genes Dev. 10.1101/gad.349917.122.

Cai, X., Gao, L., Teng, L., Ge, J., Oo, Z.M., Kumar, A.R., Gilliland, D.G., Mason, P.J., Tan, K., and Speck, N.A. (2015). Runx1 Deficiency Decreases Ribosome Biogenesis and Confers Stress Resistance to Hematopoietic Stem and Progenitor Cells. Cell Stem Cell 17, 165–177. 10.1016/j.stem.2015.06.002.

Chen, C., Liu, Y., Liu, R., Ikenoue, T., Guan, K.L., Liu, Y., and Zheng, P. (2008a). TSC-mTOR maintains quiescence and function of hematopoietic stem cells by repressing mitochondrial biogenesis and reactive oxygen species. J Exp Med 205, 2397–2408. 10.1084/jem.20081297.

Chen, C., Liu, Y., Liu, Y., and Zheng, P. (2009). mTOR regulation and therapeutic rejuvenation of aging hematopoietic stem cells. Sci Signal 2, ra75. 10.1126/scisignal.2000559.

Chen, W., Kumar, A.R., Hudson, W.A., Li, Q., Wu, B., Staggs, R.A., Lund, E.A., Sam, T.N., and Kersey, J.H. (2008b). Malignant transformation initiated by Mll-AF9: gene dosage and critical target cells. Cancer Cell 13, 432–440. 10.1016/j.ccr.2008.03.005.

Chung, S.S., Eng, W.S., Hu, W., Khalaj, M., Garrett-Bakelman, F.E., Tavakkoli, M., Levine, R.L., Carroll, M., Klimek, V.M., Melnick, A.M., and Park, C.Y. (2017). CD99 is a therapeutic target on disease stem cells in myeloid malignancies. Sci Transl Med 9. 10.1126/scitranslmed.aaj2025.

Dobin, A., Davis, C.A., Schlesinger, F., Drenkow, J., Zaleski, C., Jha, S., Batut, P., Chaisson, M., and Gingeras, T.R. (2013). STAR: ultrafast universal RNA-seq aligner. Bioinformatics 29, 15–21. 10.1093/bioinformatics/bts635.

DuRose, J.B., Scheuner, D., Kaufman, R.J., Rothblum, L.I., and Niwa, M. (2009). Phosphorylation of eukaryotic translation initiation factor 2alpha coordinates rRNA transcription and translation inhibition during endoplasmic reticulum stress. Mol Cell Biol 29, 4295–4307. 10.1128/MCB.00260-09.

Eppert, K., Takenaka, K., Lechman, E.R., Waldron, L., Nilsson, B., van Galen, P., Metzeler, K.H., Poeppl, A., Ling, V., Beyene, J., et al. (2011). Stem cell gene expression programs influence clinical outcome in human leukemia. Nat Med 17, 1086–1093. 10.1038/nm.2415.

Goardon, N., Marchi, E., Atzberger, A., Quek, L., Schuh, A., Soneji, S., Woll, P., Mead, A., Alford, K.A., Rout, R., et al. (2011). Coexistence of LMPP-like and GMP-like leukemia stem cells in acute myeloid leukemia. Cancer Cell 19, 138–152. 10.1016/j.ccr.2010.12.012.

Goncalves, K.A., Silberstein, L., Li, S., Severe, N., Hu, M.G., Yang, H., Scadden, D.T., and Hu, G.F. (2016). Angiogenin Promotes Hematopoietic Regeneration by Dichotomously Regulating Quiescence of Stem and Progenitor Cells. Cell 166, 894–906. 10.1016/j.cell.2016.06.042.

Hatlen, M.A., Arora, K., Vacic, V., Grabowska, E.A., Liao, W., Riley-Gillis, B., Oschwald, D.M., Wang, L., Joergens, J.E., Shih, A.H., et al. (2016). Integrative genetic analysis of mouse and human AML identifies cooperating disease alleles. J Exp Med 213, 25–34. 10.1084/jem.20150524.

Hidalgo San Jose, L., and Signer, R.A.J. (2019). Cell-type-specific quantification of protein synthesis in vivo. Nat Protoc 14, 441–460. 10.1038/s41596-018-0100-z.

Hidalgo San Jose, L., Sunshine, M.J., Dillingham, C.H., Chua, B.A., Kruta, M., Hong, Y., Hatters, D.M., and Signer, R.A.J. (2020). Modest Declines in Proteome Quality Impair Hematopoietic Stem Cell Self-Renewal. Cell Rep 30, 69–80 e66. 10.1016/j.celrep.2019.12.003.

Ingolia, N.T. (2016). Ribosome Footprint Profiling of Translation throughout the Genome. Cell 165, 22–33. 10.1016/j.cell.2016.02.066.

Ingolia, N.T., Brar, G.A., Rouskin, S., McGeachy, A.M., and Weissman, J.S. (2012). The ribosome profiling strategy for monitoring translation in vivo by deep sequencing of ribosome-protected mRNA fragments. Nat Protoc 7, 1534–1550. 10.1038/nprot.2012.086.

Jaako, P., Flygare, J., Olsson, K., Quere, R., Ehinger, M., Henson, A., Ellis, S., Schambach, A., Baum, C., Richter, J., et al. (2011). Mice with ribosomal protein S19 deficiency develop bone marrow failure and symptoms like patients with Diamond-Blackfan anemia. Blood 118, 6087–6096. 10.1182/blood-2011-08-371963.

Jamieson, C.H., Ailles, L.E., Dylla, S.J., Muijtjens, M., Jones, C., Zehnder, J.L., Gotlib, J., Li, K., Manz, M.G., Keating, A., et al. (2004). Granulocyte-macrophage progenitors as candidate leukemic stem cells in blast-crisis CML. N Engl J Med 351, 657–667. 10.1056/NEJMoa040258.

Jan, M., Snyder, T.M., Corces-Zimmerman, M.R., Vyas, P., Weissman, I.L., Quake, S.R., and Majeti, R. (2012). Clonal evolution of preleukemic hematopoietic stem cells precedes human acute myeloid leukemia. Sci Transl Med 4, 149ra118. 10.1126/scitranslmed.3004315.

Kharas, M.G., Okabe, R., Ganis, J.J., Gozo, M., Khandan, T., Paktinat, M., Gilliland, D.G., and Gritsman, K. (2010). Constitutively active AKT depletes hematopoietic stem cells and induces leukemia in mice. Blood 115, 1406–1415. 10.1182/blood-2009-06-229443.

Kimball, S.R., Fabian, J.R., Pavitt, G.D., Hinnebusch, A.G., and Jefferson, L.S. (1998). Regulation of Guanine Nucleotide Exchange through Phosphorylation of Eukaryotic Initiation Factor eIF2α: ROLE OF THE α-AND δ-SUBUNITS OF eIF2B*. Journal of Biological Chemistry 273, 12841–12845. https://doi.org/10.1074/jbc.273.21.12841.

Krishnamoorthy, T., Pavitt, G.D., Zhang, F., Dever, T.E., and Hinnebusch, A.G. (2001). Tight Binding of the Phosphorylated – Subunit of Initiation Factor 2 (eIF2–) to the Regulatory Subunits of Guanine Nucleotide Exchange Factor eIF2B Is Required for Inhibition of Translation Initiation. Molecular and Cellular Biology 21, 5018–5030. doi:10.1128/MCB.21.15.5018-5030.2001.

Krivtsov, A.V., Twomey, D., Feng, Z., Stubbs, M.C., Wang, Y., Faber, J., Levine, J.E., Wang, J., Hahn, W.C., Gilliland, D.G., et al. (2006). Transformation from committed progenitor to leukaemia stem cell initiated by MLL-AF9. Nature 442, 818–822. nature04980 [pii] 10.1038/nature04980.

Kruta, M., Sunshine, M.J., Chua, B.A., Fu, Y., Chawla, A., Dillingham, C.H., Hidalgo San Jose, L., De Jong, B., Zhou, F.J., and Signer, R.A.J. (2021). Hsf1 promotes hematopoietic stem cell fitness and proteostasis in response to ex vivo culture stress and aging. Cell Stem Cell 28, 1950–1965 e1956. 10.1016/j.stem.2021.07.009.

Lapidot, T., Sirard, C., Vormoor, J., Murdoch, B., Hoang, T., Caceres-Cortes, J., Minden, M., Paterson, B., Caligiuri, M.A., and Dick, J.E. (1994). A cell initiating human acute myeloid leukaemia after transplantation into SCID mice. Nature 367, 645–648. 10.1038/367645a0.

Lee, J.Y., Nakada, D., Yilmaz, O.H., Tothova, Z., Joseph, N.M., Lim, M.S., Gilliland, D.G., and Morrison, S.J. (2010). mTOR activation induces tumor suppressors that inhibit leukemogenesis and deplete hematopoietic stem cells after Pten deletion. Cell Stem Cell 7, 593–605. 10.1016/j.stem.2010.09.015.

Liu, J., Xu, Y., Stoleru, D., and Salic, A. (2012). Imaging protein synthesis in cells and tissues with an alkyne analog of puromycin. Proc Natl Acad Sci U S A 109, 413–418. 10.1073/pnas.1111561108.

Llorens-Bobadilla, E., Zhao, S., Baser, A., Saiz-Castro, G., Zwadlo, K., and Martin-Villalba, A. (2015). Single-Cell Transcriptomics Reveals a Population of Dormant Neural Stem Cells that Become Activated upon Brain Injury. Cell Stem Cell 17, 329–340. 10.1016/j.stem.2015.07.002.

Love, M.I., Huber, W., and Anders, S. (2014). Moderated estimation of fold change and dispersion for RNA-seq data with DESeq2. Genome Biol 15, 550. 10.1186/s13059-014-0550-8.

Magee, J.A., Ikenoue, T., Nakada, D., Lee, J.Y., Guan, K.L., and Morrison, S.J. (2012). Temporal changes in PTEN and mTORC2 regulation of hematopoietic stem cell self-renewal and leukemia suppression. Cell Stem Cell 11, 415–428. 10.1016/j.stem.2012.05.026.

Majeti, R., Becker, M.W., Tian, Q., Lee, T.L., Yan, X., Liu, R., Chiang, J.H., Hood, L., Clarke, M.F., and Weissman, I.L. (2009). Dysregulated gene expression networks in human acute myelogenous leukemia stem cells. Proc Natl Acad Sci U S A 106, 3396–3401. 0900089106 [pii] 10.1073/pnas.0900089106.

McGowan, K.A., Pang, W.W., Bhardwaj, R., Perez, M.G., Pluvinage, J.V., Glader, B.E., Malek, R., Mendrysa, S.M., Weissman, I.L., Park, C.Y., and Barsh, G.S. (2011). Reduced ribosomal protein gene dosage and p53 activation in low-risk myelodysplastic syndrome. Blood 118, 3622–3633. 10.1182/blood-2010-11-318584.

McKeown, M.R., Corces, M.R., Eaton, M.L., Fiore, C., Lee, E., Lopez, J.T., Chen, M.W., Smith, D., Chan, S.M., Koenig, J.L., et al. (2017). Superenhancer Analysis Defines Novel Epigenomic Subtypes of Non-APL AML, Including an RARalpha Dependency Targetable by SY-1425, a Potent and Selective RARalpha Agonist. Cancer Discov 7, 1136–1153. 10.1158/2159-8290.CD-17-0399.

Morrison, S.J., Wright, D.E., and Weissman, I.L. (1997). Cyclophosphamide/granulocyte colony-stimulating factor induces hematopoietic stem cells to proliferate prior to mobilization. Proc Natl Acad Sci U S A 94, 1908–1913. 10.1073/pnas.94.5.1908.

Pabst, C., Bergeron, A., Lavallee, V.P., Yeh, J., Gendron, P., Norddahl, G.L., Krosl, J., Boivin, I., Deneault, E., Simard, J., et al. (2016). GPR56 identifies primary human acute myeloid leukemia cells with high repopulating potential in vivo. Blood 127, 2018–2027. 10.1182/blood-2015-11-683649.

Palchaudhuri, R., Saez, B., Hoggatt, J., Schajnovitz, A., Sykes, D.B., Tate, T.A., Czechowicz, A., Kfoury, Y., Ruchika, F., Rossi, D.J., et al. (2016). Non-genotoxic conditioning for hematopoietic stem cell transplantation using a hematopoietic-cell-specific internalizing immunotoxin. Nat Biotechnol 34, 738–745. 10.1038/nbt.3584.

Papaemmanuil, E., Gerstung, M., Bullinger, L., Gaidzik, V.I., Paschka, P., Roberts, N.D., Potter, N.E., Heuser, M., Thol, F., Bolli, N., et al. (2016). Genomic Classification and Prognosis in Acute Myeloid Leukemia. N Engl J Med 374, 2209–2221. 10.1056/NEJMoa1516192.

Park, H.J., Byun, D., Lee, A.H., Kim, J.H., Ban, Y.L., Araki, M., Araki, K., Yamamura, K., Kim, I., Park, S.H., and Jung, K.C. (2012). CD99-dependent expansion of myeloid-derived suppressor cells and attenuation of graft-versus-host disease. Mol Cells 33, 259–267. 10.1007/s10059-012-2227-z.

Raffel, S., Klimmeck, D., Falcone, M., Demir, A., Pouya, A., Zeisberger, P., Lutz, C., Tinelli, M., Bischel, O., Bullinger, L., et al. (2020). Quantitative proteomics reveals specific metabolic features of acute myeloid leukemia stem cells. Blood 136, 1507–1519. 10.1182/blood.2019003654.

Rao, S., Lee, S.Y., Gutierrez, A., Perrigoue, J., Thapa, R.J., Tu, Z., Jeffers, J.R., Rhodes, M., Anderson, S., Oravecz, T., et al. (2012). Inactivation of ribosomal protein L22 promotes transformation by induction of the stemness factor, Lin28B. Blood 120, 3764–3773. 10.1182/blood-2012-03-415349.

Recher, C., Beyne-Rauzy, O., Demur, C., Chicanne, G., Dos Santos, C., Mas, V.M., Benzaquen, D., Laurent, G., Huguet, F., and Payrastre, B. (2005). Antileukemic activity of rapamycin in acute myeloid leukemia. Blood 105, 2527–2534. 10.1182/blood-2004-06-2494.

Sampath, P., Pritchard, D.K., Pabon, L., Reinecke, H., Schwartz, S.M., Morris, D.R., and Murry, C.E. (2008). A hierarchical network controls protein translation during murine embryonic stem cell self-renewal and differentiation. Cell Stem Cell 2, 448–460. 10.1016/j.stem.2008.03.013.

Sanchez, C.G., Teixeira, F.K., Czech, B., Preall, J.B., Zamparini, A.L., Seifert, J.R., Malone, C.D., Hannon, G.J., and Lehmann, R. (2016). Regulation of Ribosome Biogenesis and Protein Synthesis Controls Germline Stem Cell Differentiation. Cell Stem Cell 18, 276–290. 10.1016/j.stem.2015.11.004.

Shlush, L.I., Zandi, S., Mitchell, A., Chen, W.C., Brandwein, J.M., Gupta, V., Kennedy, J.A., Schimmer, A.D., Schuh, A.C., Yee, K.W., et al. (2014). Identification of pre-leukaemic haematopoietic stem cells in acute leukaemia. Nature 506, 328–333. 10.1038/nature13038.

Signer, R.A., Magee, J.A., Salic, A., and Morrison, S.J. (2014). Haematopoietic stem cells require a highly regulated protein synthesis rate. Nature 509, 49–54. 10.1038/nature13035.

Signer, R.A., Qi, L., Zhao, Z., Thompson, D., Sigova, A.A., Fan, Z.P., DeMartino, G.N., Young, R.A., Sonenberg, N., and Morrison, S.J. (2016). The rate of protein synthesis in hematopoietic stem cells is limited partly by 4E-BPs. Genes Dev 30, 1698–1703. 10.1101/gad.282756.116.

Somervaille, T.C., and Cleary, M.L. (2006). Identification and characterization of leukemia stem cells in murine MLL-AF9 acute myeloid leukemia. Cancer Cell 10, 257–268. 10.1016/j.ccr.2006.08.020.

Subramanian, A., Tamayo, P., Mootha, V.K., Mukherjee, S., Ebert, B.L., Gillette, M.A., Paulovich, A., Pomeroy, S.L., Golub, T.R., Lander, E.S., and Mesirov, J.P. (2005). Gene set enrichment analysis: a knowledge-based approach for interpreting genome-wide expression profiles. Proc Natl Acad Sci U S A 102, 15545–15550. 10.1073/pnas.0506580102.

Vaikari, V.P., Du, Y., Wu, S., Zhang, T., Metzeler, K., Batcha, A.M.N., Herold, T., Hiddemann, W., Akhtari, M., and Alachkar, H. (2020). Clinical and preclinical characterization of CD99 isoforms in acute myeloid leukemia. Haematologica 105, 999–1012. 10.3324/haematol.2018.207001.

van Galen, P., Kreso, A., Mbong, N., Kent, D.G., Fitzmaurice, T., Chambers, J.E., Xie, S., Laurenti, E., Hermans, K., Eppert, K., et al. (2014). The unfolded protein response governs integrity of the haematopoietic stem-cell pool during stress. Nature 510, 268–272. 10.1038/nature13228.

van Galen, P., Mbong, N., Kreso, A., Schoof, E.M., Wagenblast, E., Ng, S.W.K., Krivdova, G., Jin, L., Nakauchi, H., and Dick, J.E. (2018). Integrated Stress Response Activity Marks Stem Cells in Normal Hematopoiesis and Leukemia. Cell Rep 25, 1109–1117 e1105. 10.1016/j.celrep.2018.10.021.

Wang, F., Morita, K., DiNardo, C.D., Furudate, K., Tanaka, T., Yan, Y., Patel, K.P., MacBeth, K.J., Wu, B., Liu, G., et al. (2021a). Leukemia stemness and co-occurring mutations drive resistance to IDH inhibitors in acute myeloid leukemia. Nat Commun 12, 2607. 10.1038/s41467-021-22874-x.

Wang, V.E., Blaser, B.W., Patel, R.K., Behbehani, G.K., Rao, A.A., Durbin-Johnson, B., Jiang, T., Logan, A.C., Settles, M., Mannis, G.N., et al. (2021b). Inhibition of MET Signaling with Ficlatuzumab in Combination with Chemotherapy in Refractory AML: Clinical Outcomes and High-Dimensional Analysis. Blood Cancer Discovery. 10.1158/2643-3230.Bcd-21-0055.

Yan, M., Kanbe, E., Peterson, L.F., Boyapati, A., Miao, Y., Wang, Y., Chen, I.M., Chen, Z., Rowley, J.D., Willman, C.L., and Zhang, D.E. (2006). A previously unidentified alternatively spliced isoform of t(8;21) transcript promotes leukemogenesis. Nat Med 12, 945–949. 10.1038/nm1443.

Yilmaz, O.H., Valdez, R., Theisen, B.K., Guo, W., Ferguson, D.O., Wu, H., and Morrison, S.J. (2006). Pten dependence distinguishes haematopoietic stem cells from leukaemia-initiating cells. Nature 441, 475–482. nature04703 [pii] 10.1038/nature04703.

Zhang, J., Grindley, J.C., Yin, T., Jayasinghe, S., He, X.C., Ross, J.T., Haug, J.S., Rupp, D., Porter-Westpfahl, K.S., Wiedemann, L.M., et al. (2006). PTEN maintains haematopoietic stem cells and acts in lineage choice and leukaemia prevention. Nature 441, 518–522. nature04747 [pii] 10.1038/nature04747.

Zismanov, V., Chichkov, V., Colangelo, V., Jamet, S., Wang, S., Syme, A., Koromilas, A.E., and Crist, C. (2016). Phosphorylation of eIF2alpha Is a Translational Control Mechanism Regulating Muscle Stem Cell Quiescence and Self-Renewal. Cell Stem Cell 18, 79–90. 10.1016/j.stem.2015.09.020.

